# Paternal cardiac injury elicits an inflammatory signal relay to the gonads with intergenerational cardiac effects in vertebrates

**DOI:** 10.64898/2026.08.22.746193

**Authors:** Benedetta Coppe, Prateek Arora, María Galardi-Castilla, Andrés Sanz-Morejón, Théo Meister, Ksenia Skvortsova, Barbara Kupferschmid, Ayisha Marwa Mangattu Parambil, Nick Kirschke, Gianrico Gadient, Ines J. Marques, Emrush Rexhaj, Ozren Bogdanovic, Nadia Mercader

## Abstract

The blood-gonadal barrier protects the germline from parental exposures. A phenomenon known as intergenerational inheritance suggests that, exceptionally, this barrier can be surpassed with consequences for the subsequent generation. Specific diet regimes and early traumatic experiences have been among the chronic stressors shown to be able to lead to intergenerational inheritance in mammals. Less is known about how acute stress can affect the germline.

Cardiac damage leads to several alterations in peripheral organs and, overall, affects blood flow, metabolism, and the immune response. Whether cardiac damage can also affect the reproductive system is not known and might offer new insights into the potential inheritance of cardiovascular disease.

Here, we used zebrafish and mouse models to explore the intergenerational role of cardiac damage and repair. In the first week after a cardiac cryolesion, male zebrafish gonads and gametes activated responses associated with inflammation. In sperm, chromatin accessibility was found altered in response to cardiac cryolesion. Offspring of cryoinjured zebrafish males revealed changes in cardiac function and cardiac gene expression. Induction of systemic sterile inflammation in the paternal generation mimicked cardiac injury effects in the following generation, while anti-inflammatory treatments in the injured paternal generation partially recovered F1 cardiac features. Similar features were found in mouse testis after a neonatal injury, and in the hearts of their offspring, suggesting a conserved role of sterile inflammation as a vector for intergenerational transmission of cardiac injury.

**Teaser:** A cardiac injury triggers inflammation. This affects the parental germline and leads to phenotypic alterations in offspring.

**Graphical abstract:** 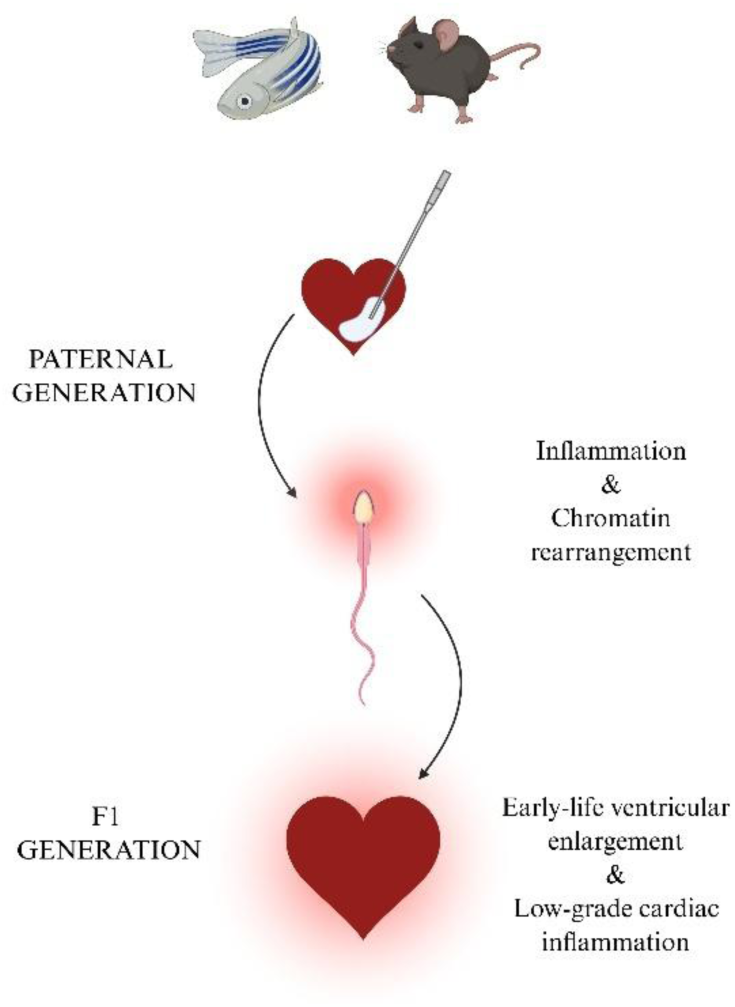

## INTRODUCTION

Exposure to environmental stressors, such as chemicals, diet, and trauma, alters gene regulation in somatic cells and germline, and several reports suggest that these have consequences for the next generation. Epigenetic modifications of the gametes are a vector for transmission of the experienced stressor from the parental generation to their offspring. Epigenetic regulation of gene expression occurs at the chromatin level, *via* regulation of chromatin accessibility through histone post-translational modifications (PTMs) and DNA methylation, or by gene silencing through activation of small RNA silencing pathways. These epigenetic pathways can be altered in the germline of the exposed generation in response to different *stimuli,* and such alterations can correlate with the establishment of specific phenotypic manifestations in the following generations. This mechanism is known as epigenetic inheritance (Fitz-James & Cavalli, 2022; Skvortsova et al., 2018).

One of the best-established stressors leading to epigenetic inheritance is the diet regime. Chronic episodes of starvation or prolonged high-fat diet exposure can lead to metabolic disorders through the *patriline* in different organisms, ranging from C. elegans and Drosophila to mice and humans, as evidenced by experimental and epidemiological data (Carone et al., 2010; Guida et al., 2019; Rechavi et al., 2014; Tomar et al., 2024). Less is known about the effect an acute stressor, such as an organ injury, can have on the next generation. There is evidence that upon liver damage, DNA methylation is altered in the gametes (Zeybel et al., 2012); however, phenotypic alterations in the next generations are debated (Beil et al., 2023).

Cardiovascular diseases (CVD), and among them, heart failure, are systemic disorders characterized by heart-to-peripheral organs bidirectional communication involving immune, metabolic, neural, endocrine, and mechanical signals (Schiattarella & Kontaridis, 2025). Cardiac injury elicits the activation of the immune system in several peripheral tissues in different animal models and in humans (Hoyer et al., 2019; Cortada et al., 2024; Rettkowski et al., 2025). It remains unexplored whether interactions between the heart and the reproductive system occur, possibly influencing information carried by the gametes. In humans, a cohort study performed on adults with CVD identified an increased risk of developing CVD in children during childhood and young adulthood. Among risk factors influencing the cardiovascular development of offspring are poor uterine vascularization, genetic predisposition, and pre- and postnatal influence of family lifestyle (Hossin et al., 2024). Indeed, nutrition, smoking, and alcohol consumption have been associated with a higher risk of cardiac malformations in offspring (Peng et al., 2019). It remains unexplored whether myocardial infarction (MI) experienced in adulthood can alone influence gametes and the following generation. Given that the prevalence of an MI episode can encompass the reproductive age span (Ramphul et al., 2021), it is of great interest to explore its possible consequences for the next generation.

To understand the intergenerational effects of MI, we used the mouse and zebrafish experimental models that allow us to control other previously mentioned confounding exposures. We then evaluated potential changes in gonadal gene and protein expression, together with molecular alterations at the spermatozoal level. We also assessed the effect of paternal cryolesion on the subsequent generation, focusing in particular on the effect on cardiac function. Overall, our results suggest that cardiac damage leads to inflammation and alters spermatogenesis progression in parental male gonads as well as chromatin accessibility in mature gametes. The alterations in the parental gametes correlate with changes in cardiac morphology and function in offspring, both in zebrafish and mice.

Overall, our work suggests that a cardiac injury triggers sterile inflammation that can affect the germline and lead to alterations in cardiac function in offspring.

## RESULTS

### Cardiac injury activates an immune response in zebrafish male gonads

Zebrafish can regenerate their hearts after severe cardiac injury (Poss et al., 2002; González-Rosa et al., 2011). The model is therefore suited to start exploring effects of a cardiac lesion on the next generation. To investigate the consequences of cardiac injury on the reproductive system, we collected wild-type zebrafish testes from uninjured males or males at 1, 7, or 133 days after cardiac cryoinjury (CI) and performed poly(A) bulk RNA-seq at all time points (Figure 1Aa).

To minimize batch effect, testes were collected at the same time, while the injury was performed on different days before euthanizing the animals. Control animals (*i.e.,* “uninjured”) were anesthetized 7 days before testes extraction but did not undergo sham surgery.

Principal component analysis (PCA) and heatmap of differentially expressed genes (DEGs), defined by padj ≤ 0.05 and |Log_2_FoldChange| ≥ 1, showed that the expression profile of samples collected at 7 days post-injury (dpi) differed the most compared to the others (Figure S1Aa and Ab), suggesting a transient response to cardiac CI in the gonads. An initial modest gene expression response was detected at 1 dpi (39 DEGs), which became more pronounced at 7 dpi (210 DEGs), and subsequently declined at 133 dpi (34 DEGs) (Figures 1Ab,Ac, Figures S1Ba,b, and Bulk_RNA-seq_testes_ZF).

Gene Ontology (GO) biological processes analysis of DEGs at 7 dpi revealed significant enrichment of pathways involved in inflammation, among others (Figure 1Ad, Figure S1C, and Bulk_RNA-seq_testes_ZF). DEGs taking part in inflammation-associated pathways (i.e. “regulation of inflammatory response” and “acute inflammatory response”) and found upregulated at 7 dpi were *Interferon Lambda Receptor 1* (*ifnlr1*), *NLR Family Pyrin Domain Containing 5* (*nlrp5*), *Solute Carrier Family 7 Member 2 (slc7a2), alpha-2-HS-glycoprotein–1 (ahsg1),* and *inter-alpha-trypsin inhibitor heavy chain 3b, tandem duplicate 2 (itih3b.2)*. Within the same GO terms, we observed the downregulation of *nuclear factor of kappa light polypeptide gene enhancer in B-cells inhibitor, alpha b (Nfkbiab), suppressor of cytokine signaling 3a (socs3*a), *CCAAT enhancer binding protein beta (cebpb), ghrelin/obestatin prepropeptide (ghrl),* and *serum amyloid A (saa)*. Of note, *soc3a, cebpb, and ifnlr1* are associated with the pro-inflammatory JAK/STAT pathway (Elsaeidi et al., 2014; Qin et al., 2021; Ren et al., 2023), while *nfkbiab and ghrl* are implicated in the negative regulation of *NF-kB* (Nuclear Factor-kappa B) signal transduction (Correa et al., 2004; W. G. Li et al., 2004).

Among the downregulated genes associated with the negative regulation of phosphorus/phosphate metabolic processes were *DNA-damage-inducible transcript 4 (ddit4*), which inhibits mTOR signalling to control cell growth in response to stress (Sofer et al., 2005); *Suppressor Of Cytokine Signaling 3 (socs3*), which negatively regulates JAK/STAT-mediated cytokine pathways (Y. Yin et al., 2015); *Dual Specificity Phosphatase 6 (dusp6*), which attenuates MAPK/ERK pathway activity (Furukawa et al., 2008; Muda et al., 1996); and the two paralogs *growth arrest and DNA-damage-inducible, gamma a and b tandem duplicate 1 (gadd45ga* and *gadd45gb*.1), which are involved in cell-cycle regulation and stress-induced growth arrest (Jin et al., 2000) (Figure 1Ad and Figure S1Ca,b). Together, these genes are inovlved in the regulation of cell growth, proliferation, and inflammatory signaling pathways through pathways including mTOR, JAK/STAT, and MAP/ERK. In addition to these genes, among the top DEGs at 7 dpi, *MYCN proto-oncogene, bHLH transcription factor (mycn),* involved in pluripotency and the regulation of spermatogonial stem cells differentiation (Reza et al., 2025), and *laminin gamma 2 (lamc2*), an extracellular matrix (ECM) component that contributes to epithelial basement membrane integrity, were highly expressed at 7 dpi. This may indicate alterations in barrier integrity and potential permeabilization with effects on spermatogenesis. Together, the dysregulation of these genes in the testes following cardiac damage suggests alterations in spermatogonial expansion, potentially affecting the balance between germ cell proliferation and differentiation in response to inflammation.

Next, we performed proteomics of testes from uninjured and injured fish at 1 and 7 dpi. Similarly to transcriptomic data, we collected testes from four males per replicate and compared them across 5 replicates per condition (Figure 1Ba, Bb). Extracted proteins were digested into peptides, separated by chromatography, ionized, and analysed by mass spectrometry. 6’602 total protein groups were identified after filtering. 263 proteins were identified as differentially abundant (DEGs; padj≤0.05 & |Log_2_FoldChange| ≥ 1) between 1 dpi and uninjured samples (Figure S2Aa), while 341 proteins were differentially abundant between 7 dpi and uninjured samples (Figure S2Ab, <u>Bulk_proteomics_testes_ZF</u>).

GO analysis performed on differentially abundant proteins identified pathways associated with RNA processes (*e.g.* RNA splicing, ncRNA processing, ribonucleoprotein complex biogenesis) and cell cycle (*e.g.* positive regulation of cell cycle, sister chromatid segregation, TOR signaling) at both time points (Figure 1Bc, Bd, Figure S2Aa-Bc), suggesting once again alterations in the regulation of spermatogenesis (<u>Bulk_proteomics_testes_ZF</u>). At 1 dpi, pathways associated with chromosome segregation were downregulated (Figure S2Ab), while mitotic cell cycle phase transition and mRNA processing pathways were upregulated (Figure S2Ac), possibly indicating the activation of spermatogonia-related programs. At 7 dpi, genes linked to rRNA processing and chromosome segregation were downregulated (Figure S2Bb), whereas pathways associated with mRNA processing, TORC1 signaling, and epithelial tube morphogenesis were upregulated (Figure S2Bc), indicating a shift toward post-mitotic differentiation and cyst maturation. Indeed, Hematoxylin-Eosin staining of the testis suggested that the proportion of spermatocyte-containing cysts slightly decreased, accompanied by an increase in cysts containing spermatogonia and spermatids after one week from the cardiac injury (Figure S3A,B).

Together, these molecular and morphological data indicate that a cardiac injury tightly regulates spermatogenesis in response to inflammation, possibly as an adaptive mechanism to preserve fertility.

### Cardiac injury alters chromatin accessibility in zebrafish sperm

Environmental exposures have been associated with alterations in epigenetic marks in the germline, including changes in DNA methylation (de Castro Barbosa et al., 2016; Jung et al., 2022), histone post-translational modifications (PTMs) (Ciabrelli et al., 2017; Klosin et al., 2017; Seong et al., 2011), and the accumulation of specific small non-coding RNA species (sncRNAs) (Sharma et al., 2016; Toker et al., 2022; Tomar et al., 2024b).

We assessed whether cardiac damage influences gametes chromatin accessibility through omics approaches. Changes in DNA methylation patterns were tested by Whole Genome Bisulfite sequencing (WGBS), and chromatin accessibility alterations by the Assay for Transposase-Accessible Chromatin with high-throughput sequencing (ATAC-seq).

Spermatozoa were collected from uninjured and injured males at 7 dpi by gentle suction using a capillary tube, as previously described in protocols for semen cryopreservation and *in vitro* fertilization (Carmichael et al., 2009). Sperm purity was assessed by visual inspection under the microscope (Figure S4A, Supplementary Movie S1).

Since in teleost species paternal DNA methylation patterning is largely maintained after fertilization and is matched by the maternal genome by the time of zygotic genome activation (occurring at 1 k-cells stage in the zebrafish) (Jiang et al., 2013; Potok et al., 2013; Ross et al., 2023), we first assessed changes in DNA methylation in sperm and 1-k cells embryos derived from uninjured and 7 dpi males. No differentially methylated regions (DMRs) were detected between uninjured and 7 dpi sperm samples, and only 2 regions showed increased methylation in embryos derived from injured fathers (Figure S4B-D, <u>WGBS_sperm_embryo_ZF</u>), overall suggesting DNA methylation is unlikely to contribute to intergenerational transmission of paternal cardiac damage in zebrafish.

Next, we assessed chromatin accessibility in sperm from uninjured and 7 dpi males by ATAC-seq (Figure 2A). Unlike mammals, where most of the histones are replaced by protamines, zebrafish retain most of histones in mature spermatozoa (Wu et al., 2011), which, along with their PTMs, can potentially influence gene expression and early embryonic development. Moreover, despite extensive chromatin compaction in spermatids, certain *loci* remain accessible, potentially providing a mechanism for the intergenerational transmission of gene-regulatory states (Burgos-Ruiz et al., 2026). We detected 604 regions in heterochromatin state (p value ≤ 0.05, log2FC < 0) and 569 in euchromatin state (p value ≤ 0.05, log2FC > 0) in sperm collected at 7 dpi compared to uninjured control (Figure 2B-C, ATAC-seq_sperm_ZF). Differentially accessible peaks were enriched in promoters (19.1%), exons (6.9%), introns (40.1%), and distal intergenic regions (32.1%) (Figure 2D).

**Fig. 1:**
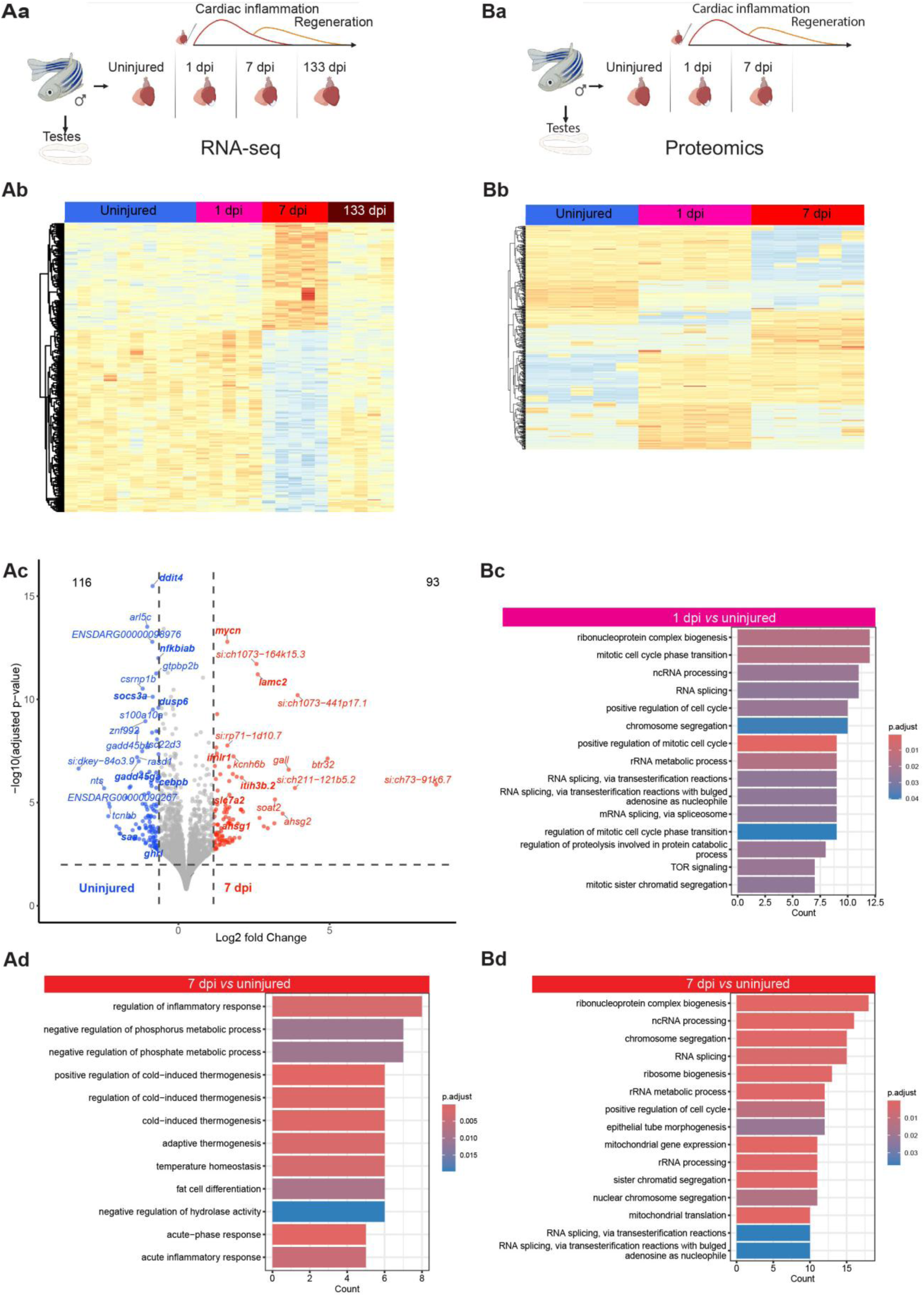
Analysis of molecular signatures in zebrafish testes following cardiac injury. **(Aa)** Schematics of the experimental design. Zebrafish males were cryoinjured and sacrificed at 1, 7, and 133 days post-injury (dpi). Transcriptomics was performed on extracted testes. Image created in BioRender. Coppe, B. (2026) https://BioRender.com/6m9279f (**Ab)** Heatmap of testes transcriptomics data. Color intensity represents gene expression level: low values (blue), high values (red) (**Ac)** Volcano plot of testes transcriptomics data at 7 dpi. The x-axis shows log₂(fold change), and the y-axis shows –log₁₀(p-value). Points in red indicate significantly upregulated genes in the injured males at 7 dpi, blue indicate significantly upregulated genes in the uninjured group, and grey indicate non-significant genes (threshold: |log₂FC| > 1, p ≤0.05). **(Ad)** GO pathway enrichment analysis - biological processes - of testes transcriptomic data (7 dpi). **(Ba)** Schematics of the experimental design. Zebrafish males were cryoinjured and euthanized at 1 and 7 dpi for protein extraction. Image created in BioRender. Coppe, B. (2026) https://BioRender.com/6m9279f (**Bb)** Heatmap of testes proteomics data. Colour intensity represents protein abundance, with blue indicating low levels and red indicating high levels, as shown in the colour bar. **(Bc)** GO Pathway enrichment analysis - biological processes - of testes transcriptomics data (1 dpi). **(Bd)** GO pathway enrichment analysis - biological processes - of testes proteomics data (7 dpi).

**Fig. 2:**
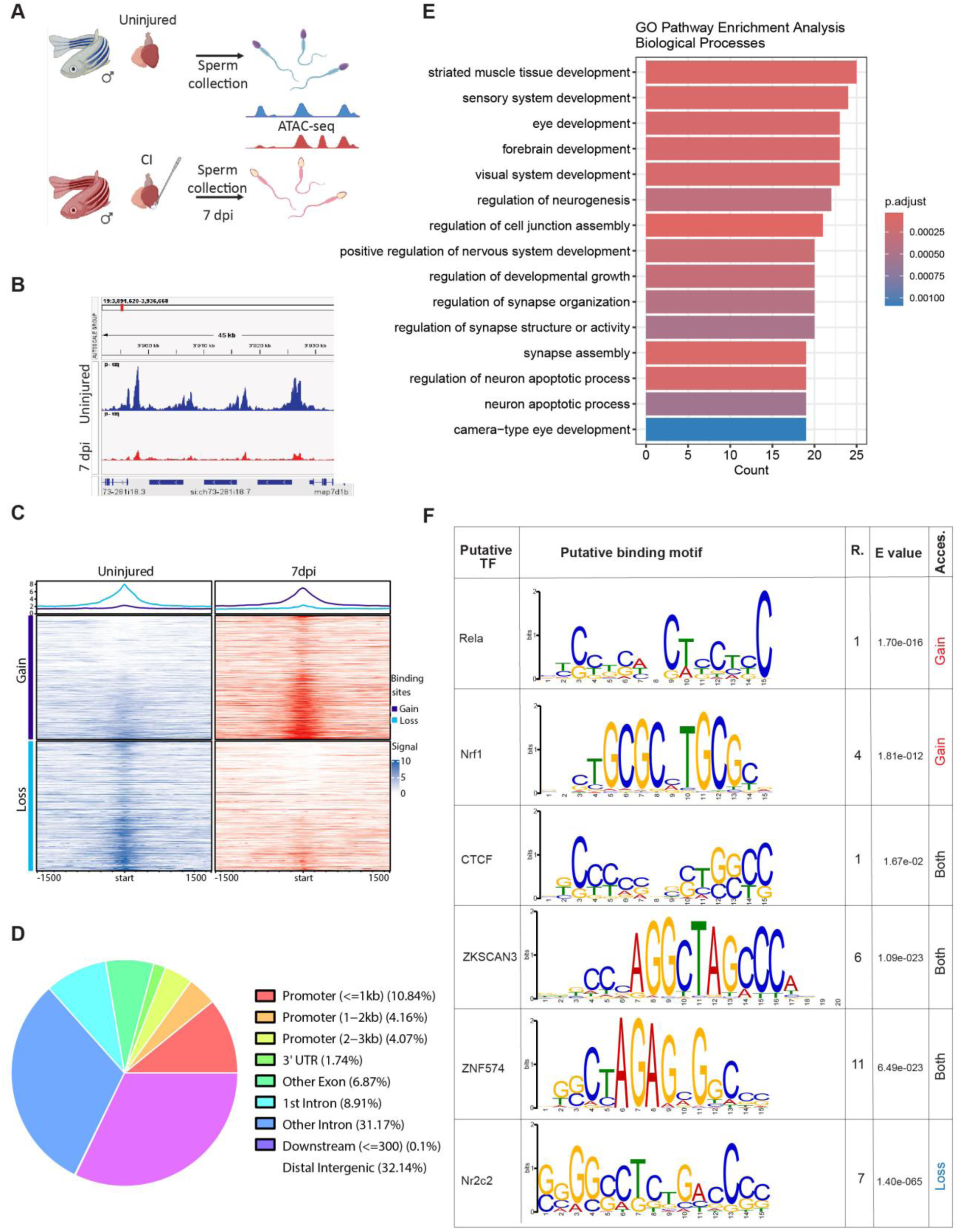
Analysis of the effect of a cardiac injury on chromatin accessibility in zebrafish sperm. **(A)** Schematics of the experimental design. Zebrafish males were cryoinjured and sperm was collected at 7 days post-injury (dpi) to perform the Assay for Transposase-Accessible Chromatin using sequencing (ATAC-seq). Sperm from uninjured control samples were collected at the same time as the injured one. Image created in BioRender. Coppe, B. (2026) https://BioRender.com/6m9279f. (**B)** IGV browser view showing an example of a differentially accessible peak detected by ATAC-seq. (**C)** Heatmaps showing signals at the peak center of differentially accessible regions. Gained and lost accessibility is shown with respect to 7 dpi samples. **(D)** Pie chart showing the genomic distribution of differential peaks between uninjured and 7 dpi samples, represented as percentages. (**E)** GO Pathway enrichment analysis - biological processes - of sperm ATAC-seq data (7 dpi). (**F)** Xstreme enrichment analysis of putative transcription binding sites in open and closed ATAC-seq regions in sperm at 7 dpi. Shown are binding motifs, R: rank (enrichment measurement of the motif based on the SEA enrichment *p*-values), E value (measurment of the statistical significance of the motif), and acc: accessibility information (Gain = euchromatic state in 7 dpi samples; Loss = heterochromatic state in 7 dpi samples; Both = TFBS found differentially accessible in both conditions).

GO enrichment analysis identified pathways related to neuronal and developmental processes (Figure 2E, <u>ATAC-seq_sperm_ZF</u>). The enrichment of neuronal GO terms likely reflects molecular mechanisms shared between neurons and sperm (*e.g.* cell adhesion, membrane organization), which might contribute to sperm maturation and function.

We performed enrichment analysis (p.val< 0.05) of putative transcription factor binding sites (TFBS) using XSTREME, a Motif Discovery and Enrichment Analysis from Meme Suite (Grant & Bailey, 2021). Among the top candidates associated with euchromatic regions, we identified binding motifs for *RELA Proto-Oncogene, NF-KB Subunit (Rela)* involved in inflammation, cell growth, and viability (Mao et al., 2025), motifs for *Nuclear Respiratory Factor 1 (Nrf1*), a regulator of mitochondrial biogenesis, cellular growth, metabolism, and oxidative stress (Hu et al., 2022), which was previously identified as potentially transmissible to the F1 generation *via* the paternal germline (Burgos-Ruiz et al., 2026), and motifs for the chromatin insulator *CCCTC-binding factor (CTCF*) (Figure FE). Notably, *CTCF* binding sites were also enriched in heterochromatic *loci* and ranked first when differentially accessible peaks were analysed together (including both open and closed differentially accessible peaks), suggesting sperm chromatin 3D reorganization following cardiac injury (<u>ATAC-seq_sperm_ZF</u>).

Predicted binding sites for several zinc finger TFs, including the autophagy repressor *ZKSCAN3* (*Zinc Finger With KRAB And SCAN Domains 3*) (Chauhan et al., 2013) and *ZNF574 (Zinc Finger Protein 574)*, were enriched in both euchromatic and heterochromatic regions. Finally, *Nr2c2* (*Nuclear Receptor Subfamily 2 Group C Member 29*), a transcription factor required for spermatogenesis (Mu et al., 2004), was enriched in heterochromatin regions in sperm at 7 dpi, further suggesting alterations during sperm maturation in response to cardiac injury.

Together, our data suggest that chromatin accessibility, but not DNA methylation, is reorganized in response to cardiac damage in male gametes.

### Zebrafish paternal cardiac injury affects cardiac function in offspring

We next asked whether the paternal history of cardiac damage could affect the next generation’s cardiac health. To assess this, we paired uninjured and injured (7 dpi) males with uninjured females of the same age (Figure 3A). While molecular signatures in both testes and sperm suggested alterations in spermatogenesis after cardiac damage, injured fathers remained fertile and successfully sired offspring at 7 dpi.

**Fig. 3:**
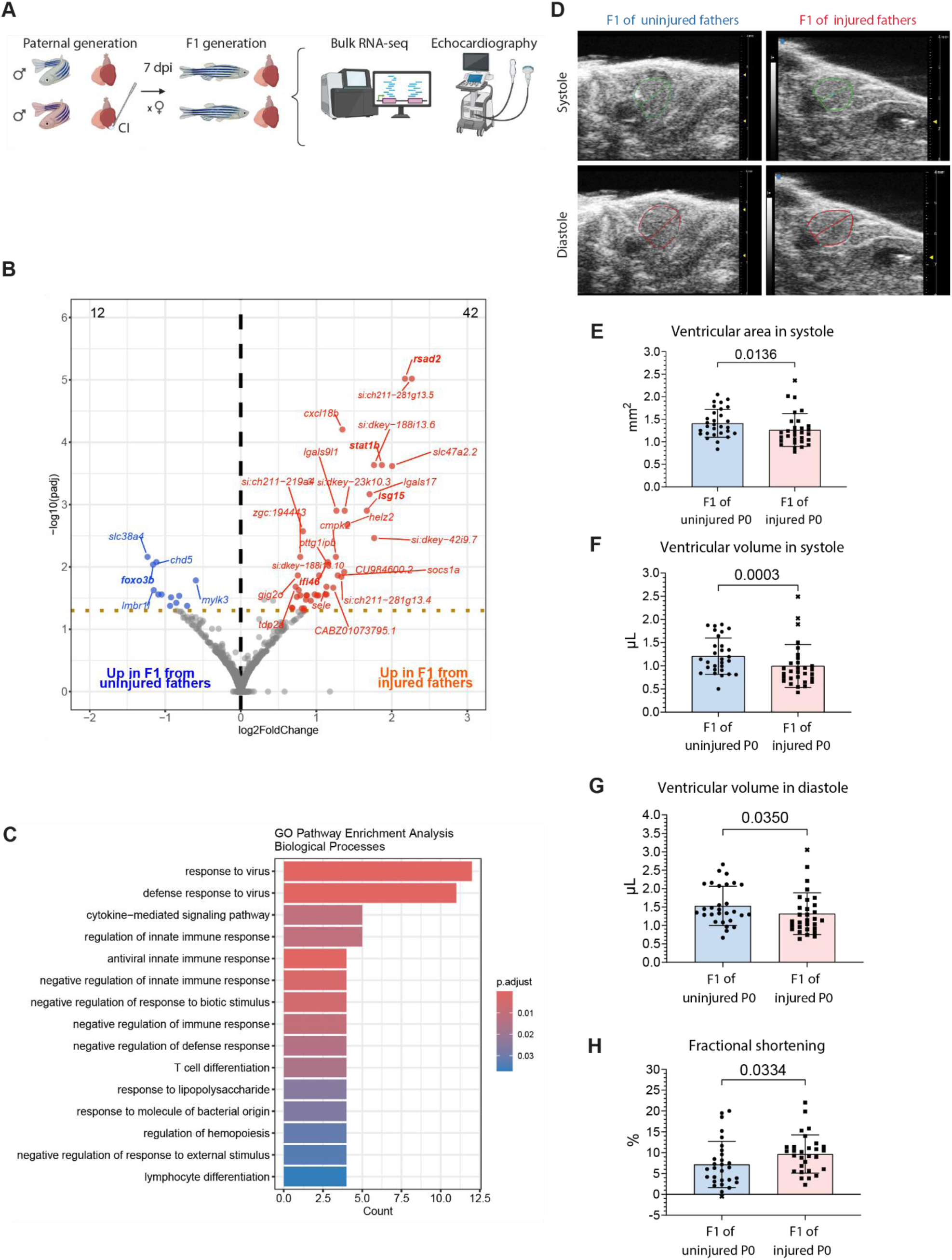
Zebrafish cardiac health assessment in the adult F1 generation of cryo-injured fathers. (**A**) Scheme of the experimental procedure. Uninjured and 7 dpi males were crossed with control females. F1 generation was grown to adulthood, when cardiac transcriptomics or echocardiographic analyses were performed in homeostatic conditions. Image created in BioRender. Coppe, B. (2026) https://BioRender.com/6m9279f (**B**) Volcano plot of genes expressed in the ventricle extracted from uninjured offspring of uninjured *vs* injured fathers (7 dpi). The x-axis shows log₂(fold change), and the y-axis shows -log₁₀(p-value). Points in red indicate significantly upregulated genes in the injured males at 7 dpi, blue indicate significantly upregulated genes in the uninjured group, and grey indicate non-significant genes (threshold: |log₂FC| > 0.58, p ≤0.05). **(C)** GO pathway enrichment analysis - biological processes - of bulk RNA-seq performed in the uninjured hearts of F1 of uninjured and 7 dpi fathers. (**D)** Echocardiography B-mode images of uninjured F1 hearts in systole (top) and diastole (bottom). (**E-H).** Quantification of echocardiography performed in the F1 generation of uninjured *vs* injured fathers. Each point corresponds to a measurement from one animal: (E) ventricular area in systole (Mann Whitney test; difference in medians, -0.1505; 95% confidence interval, -0.3215 to -0.01708), (F) ventricular volume in systole (Unpaired t test; difference in means, -0.2130; 95% confidence interval, -0.4342 to 0.008253), (G) ventricular volume in diastole (Mann Whitney test; difference in medians, -0.2221; 95% confidence interval, -0.4619 to 0.01225), (H) fractional shortening (Mann Whitney test; difference in medians, 3.545; 95% confidence interval, 0.5944 to 5.248).

As an initial approach, we aimed to understand whether paternal cardiac damage would influence the cardiac transcriptome of the next generation under physiological conditions. Hence, we performed bulk RNA-seq of ventricles extracted from the F1 generation of uninjured and injured fathers in adulthood (Figure 3A). For each replicate, we collected four hearts from siblings, with each replicate including offspring from different fathers. Differential expression analysis identified 54 genes differentially expressed (padj ≤ 0.05, |Log2FoldChange| > 0.58) between offspring of uninjured and injured fathers, of which 42 were upregulated in the offspring of injured fathers (Figure 3B, <u>Bulk_RNA-seq_F1_heart_ZF</u>). GO biological pathways enrichment analysis of DEGs identified genes such as *Radical S-adenosyl methionine domain-containing 2 (rsad2*), *signal transducer and activator of transcription 1b (stat1b), Interferon-Stimulated Gene 15 (isg15), i*nterferon-induced protein with tetratricopeptide repeats 46 (*ifi46),* and *DExH-Box Helicase 58 (dhx58*), involved in the inflammatory response, i.e. *response to virus*, *cytokine-mediated signalling pathway*, and *regulation of innate immune response* (Figure 3B,C). *Isg15* and *rsad2* are among interferon-stimulated genes commonly overexpressed upon organ injury and play key roles in propagating the inflammatory response (Carey et al., 2023; Denans et al., 2022; Wei et al., 2023.).

To assess whether the inflammation-associated signature observed in hearts from the offspring of injured males impacted cardiac morphology and functionality, we performed echocardiography in the adult F1 generation (Figure 3A). Echocardiographic analysis revealed a 10.4% reduction in ventricular area (SE = 6.0%) and a 25.1% reduction in ventricular volume (SE = 6.5%; Figure 3D, E-H) in systole in the offspring of injured *vs* uninjured fathers. Ventricular volume in diastole decreased by 17.8% (SE = 7.6%; Figure 3G), while fractional shortening increased by 29.9% (SE = 20.7%; Figure 3H). Other parameters did not show statistically significant changes (Figure S5A-E), but a tendency toward increased cardiac output (Figure S5B), ejection fraction (Figure S5C), and heart rate (Figure S5E) was observed in the offspring of injured fathers. The increased heart rate, associated with a physiological ventricular volume reduction, may be indicative of an enhanced acute stress response (Yoganathan et al., 2023) in the offspring of injured fathers. *Foxo3b* is one of the genes found to be downregulated in the offspring of injured fathers (Figure 3B). Interestingly, loss of function of *Foxo3* in mice has been shown to enhance cardiac function (Xia et al., 2025), possibly relating reduced *foxo3b* expression to the increased cardiac functionality observed in our model. Together, the partial ventricular restriction and increased cardiac contractility observed in the offspring of injured fathers under basal conditions suggest an enhanced stress response relative to controls (Yoganathan et al., 2023). Alternatively, this phenotype may represent an example of cardiac morphological restriction (Du et al., 2008) accompanied by functional compensation, potentially associated with an impaired response to damage.

Finally, we assessed whether cardiac contractility or other morphological features were already altered during development. Uninjured and injured (7 dpi) wild-type males were paired-crossed simultaneously with uninjured transgenic females expressing a fluorescent protein under the cardiomyocyte (CM) promoter *myl7* (see materials and methods). Larvae were selected based on fluorescence, and their heart imaged using a high-content fluorescent microscope at 3 dpf as previously described (Ernst et al., 2023). Quantification of functional (heart rate, cardiac output, fractional shortening) and morphological (ventricular area, volume, and circularity) cardiac features was automatically assessed using a deep learning model based on a U-Net architecture (adapted from (Ernst et al., 2023), <u>Acquifer</u>), followed by manual curation of the data (Figure 4A-C). We observed a statistically significant increment in cardiac ventricular size in the larval offspring of injured *vs* uninjured males, both in systole and diastole (*i.e.* systolic area 8.1% ± 1.9% Standard Error (SE)), and diastolic area 4.89% ± 1.3% SE) (Figure 4D-G). There was also a slight but significant increase in systolic and diastolic volume (Figure S6A,B). We further observed that fractional shortening (−4.67% ± 1.8% SE) was reduced (Figure 4H), while heart rate (6.1% ± 1.6% SE) and cardiac output (9.5% ± 3.3% SE) increased in the offspring of injured fathers (Figure 4I-J). Other parameters did not show significant differences (Figure S6C-E). When we compared body length, no changes between the two groups were observed (Figure S6F), suggesting developmental variations are likely cardiac-specific.

**Fig. 4:**
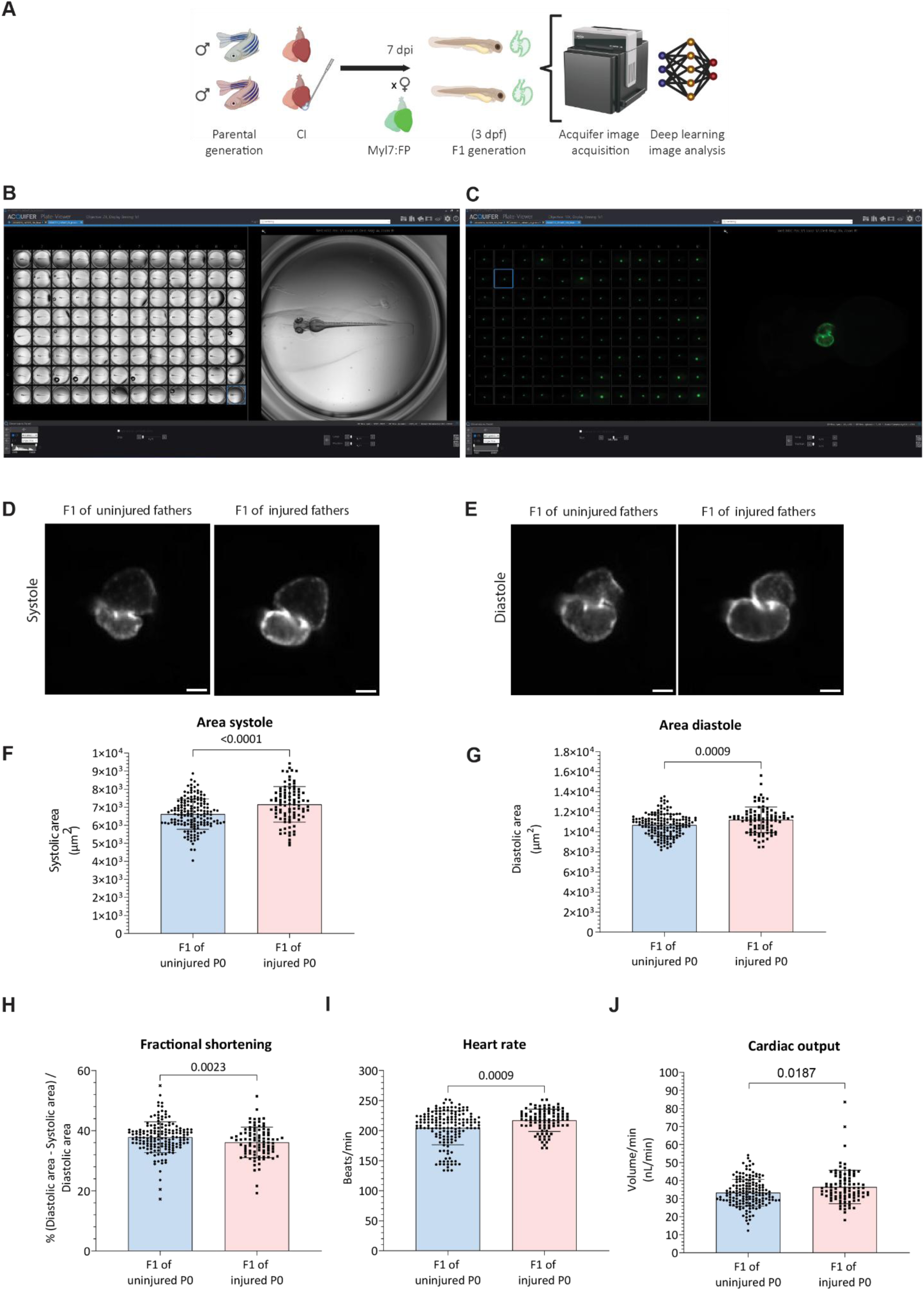
Comparison of early-life cardiac phenotype between the offspring of uninjured and 7 dpi zebrafish males. **(A)** Experimental design to investigate alterations during cardiac development in the F1 from uninjured and 7 dpi at 3 days post-fertilization (dpf). Uninjured and 7 dpi males were crossed with control transgenic females expressing a fluorescent protein under the cardiomyocytes (CMs) promoter *myl7*. F1 larvae were selected and disposed in a 96-well plate at 3 dpf. Cardiac structure and function were assessed through 300 frames using the Acquifer machine (Bruker), and image quantification was performed using a deep learning model based on a U-Net architecture. CI: cryo-injury. Image created in BioRender. Coppe, B. (2026) https://BioRender.com/6m9279f. (**B,C)** Visualization of the 96-well plate in the Plate-Viewer software of Acquifer Machine (Bruker). (B) Zebrafish embryos visualized in brightfield (2X magnification); (C). Selected hearts visualized using fluorescence (10X magnification). (**D,E)** Representative images of larval hearts from uninjured and injured males 3 dpf in systole (**D**) and diastole (**E**). Scale bar: 50 µm. (**F**) Scatter plot with SD of systolic area in the F1 of uninjured and injured males at 3 dpf (Unpaired t test; difference in means 534.5; 95% confidence interval: 303.1 to 766.0) (**G**) Scatter plot with SD of diastolic area in the F1 of uninjured and injured males at 3 dpf (Mann-Whitney test; difference in medians 513.1; 95% confidence interval: 189.2 to 759.1). (**H**) Fractional shortening in the offspring of uninjured and 7 dpi males (Unpaired t test; difference in means -1.769; 95% confidence interval: -3.081 to -0.4563). (**I)** F1 heart rate at 3 dpf (Mann-Whitney test; difference in medians 6.000; 95% confidence interval: 4.534 to 14.43). (**J**) Cardiac output measured at 3 dpf in the offspring of uninjured and injured males (Unpaired t test; difference in means 3.160; 95% confidence interval: 1.060 to 5.259).

A possible explanation for the cardiac phenotype observed at the larval stage is again an enhanced stress response, observed as increased heart rate (HR), together with enhanced development in the offspring of injured fathers, usually associated with increased ventricular dimensions, cardiac output, and reduced fractional shortening (Bagatto & Burggren, 2006; De Luca et al., 2014).

### Inflammation represents an intergenerational cardiac injury signal relay

The molecular and functional analyses of cardiac health in the offspring of injured fathers revealed low-grade immune activation and enhanced stress-related response under baseline conditions, suggesting that cardiac injury can induce inflammatory and stress alterations not only in the affected generation but also in their progeny. To investigate whether the systemic inflammatory state induced in the paternal generation following cardiac injury could be a key mediator of these intergenerational effects, we simulated a similar inflammatory state in adult males and assessed the cardiac phenotype of their offspring. Our previous molecular analyses identified altered regulation of genes involved in *NF-κB* and JAK-STAT signalling in the testis (Figure 1Ac,Ad), as well as enrichment of *RelA/NF-κB*-associated regulatory elements in sperm chromatin (Figure 2F) following cardiac damage. IFN-γ increases in the circulation after an MI (Patel et al., 2009) and is among the cytokines known to activate JAK-STAT signalling and *NF-κB*-associated inflammatory pathways (Gough et al., 2008). We therefore used the transgenic line *Tg*(*hsp70l:ifng1-2-V5,cryaa:Cerulean*), in short, hsp70:ifng1 (Sawamiphak et al., 2014), to experimentally model a systemic sterile inflammatory state in adult males and partially mimic the molecular alterations observed following cardiac injury. We performed heat shock (HS) in adult males from *hsp70:ifng1* and in control lines. All males were pair-crossed with untreated females one week later (Figure 5A). Interestingly, paternal IFN-γ systemic expression influenced F1 cardiac development similarly to a paternal cardiac injury. Cardiac ventricular dimensions were significantly larger (around 10%) in the F1 larvae from heat-shocked *hsp70:ifng1* fathers compared to the non-heat-shocked *hsp70:ifng1* controls (HS-) and heat-shocked wild-type fathers (HS+) (Figure 5B-C, Figure S7A,B). At the functional level, fractional shortening decreased by ∼3% (Figure 5D), while heart rate and cardiac output increased by around 4 and 9%, respectively (Figure 5E-F), in the offspring born following paternal IFNγ activation. Stroke volume increased in *hsp70:ifng1* groups (both HS+ and -), more pronounced in the F1 from heat-shocked *hsp70:ifng1* fathers (Figure S7C). Circularity did not differ among conditions (Figure S7E,F). Body length increased in both *hsp70:ifng1* groups, regardless of the heat shock, possibly highlighting lineage-dependent variations (Figure S7F). Taken together, these results indicate that systemic sterile inflammation in the parental generation affects cardiac development in the next generation in a similar way to cardiac damage.

**Fig. 5:**
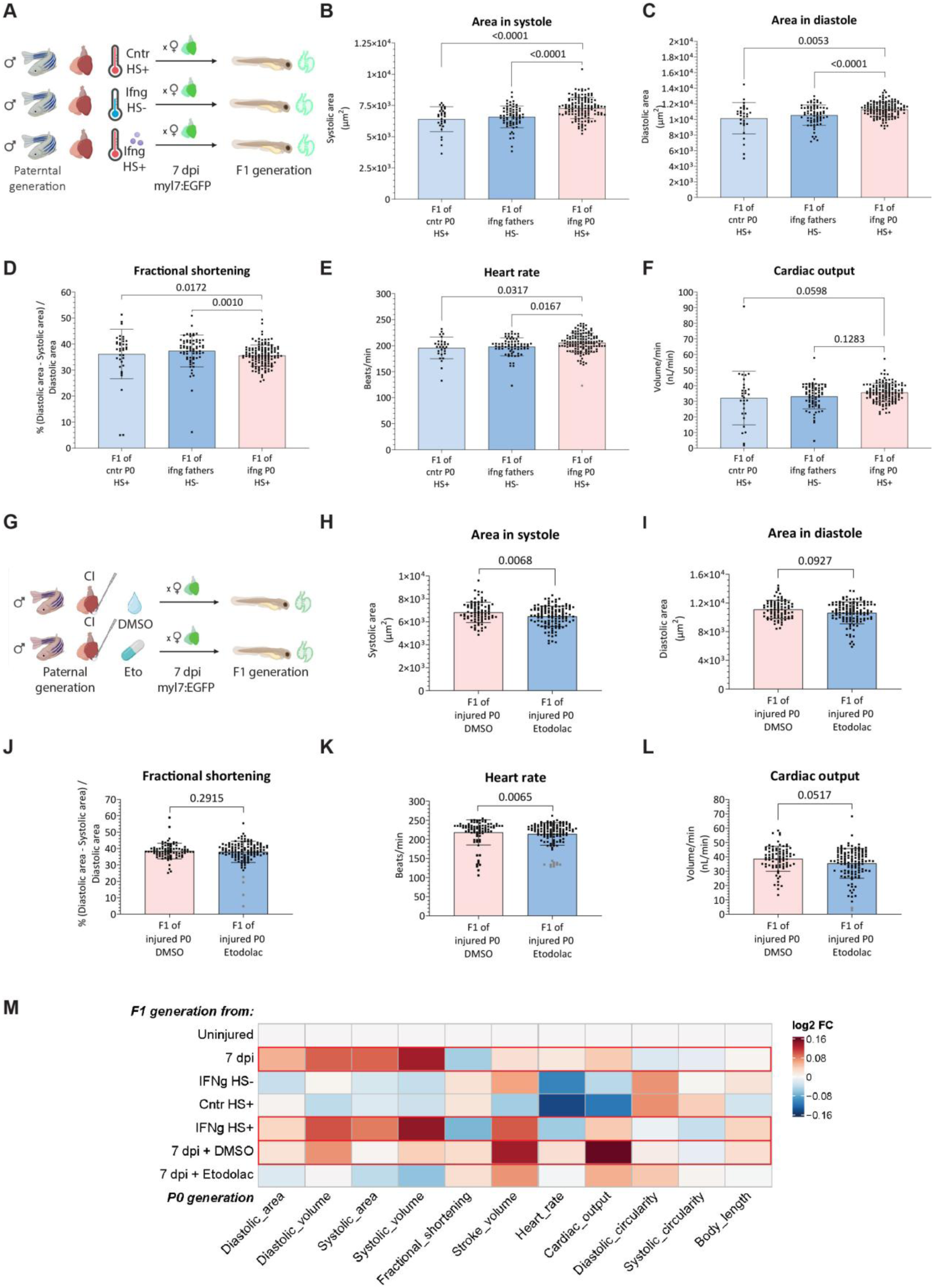
Mechanistic experiments to simulate or recover the F1 cardiac phenotype by acting on the paternal generation. **(A)** Experimental design used to simulate a systemic inflammatory response in the paternal generation by inducing IFNγ activation using the *Tg(hsp70l:ifng1-2-V5, cryaa:Cerulean)* line. Adult males underwent heat shock (HS) and were crossed one week later with untreated *Tg(myl7:EGFP)* females. Controls included transgenic males that did not undergo HS and males lacking the HS promoter that were heat shocked. Image created in BioRender. Coppe, B. (2026) https://BioRender.com/6m9279f (**B-F)**. Cardiac parameters measured at 3 dpf in the offspring of *hsp70:ifng1* (HS + and HS-) and control (HS-) fathers: B. Area in systole (One-way ANOVA, Dunn’s multiple comparisons test); **C**. Area in diastole (One-way ANOVA, Tukey’s multiple comparisons test); (**D**) Fractional shortening (One-way ANOVA, Tukey’s multiple comparisons test); (**E**) Heart rate (One-way ANOVA, Dunn’s multiple comparisons test); (**F**) Cardiac output (One-way ANOVA, Tukey’s multiple comparisons test); (**G**) Experimental design to assess the effect of paternal anti-inflammatory treatment on the next generation phenotype. Injured males (CI) were injected with 10mg/Kg Etodolac every 24 hours for 7 days, or with 1% DMSO as control. At 7 dpi, males were crossed with untreated *Tg(myl7:EGFP)* females, and cardiac development was assessed in the next generation. (**H-L**) Cardiac parameters measured at 3 dpf in the offspring of injured fathers treated with Etodolac vs control (DMSO): (**H**) Ventricular area in systole (Unpaired t test; difference in means: -348.5 ± 127.5; 95% confidence interval: -599.9 to -97.2); (**I**) Ventricular area in diastole (Mann-Whitney test; difference in medians -268.7; 95% confidence interval: -864.7 to -83.4); (**J**) Fractional shortening (Mann-Whitney test; difference in medians 0.7599; 95% confidence interval: -0.5 to 1.9; (**K)**. Heart rate (Mann- Whitney test; difference in medians -5.280; 95% confidence interval: -12.0 to -2.2); (**L**) Cardiac output (Mann-Whitney test; difference in medians -2.375; 95% confidence interval: -5.3 to -0.1). (**M**) Overview heatmap of cardiac parameters and body length measured at 3 dpf in offspring of uninjured and injured fathers, with or without treatments (systemic IFNγ activation, heat shock, Etodolac, or DMSO injection). The average value of each treatment is normalized by the control average (uninjured fathers). Morphological parameters are normalized by the body length.

Next, we wondered whether treating the parental generation with anti-inflammatory drugs would recover the phenotype in offspring. First, we validated the anti-inflammatory effect of selected anti-inflammatory compounds (*i.e*., Sulfasalazine (S. Kim et al., 2015) and Etodolac (Hall et al., 2014)) in zebrafish by assessing the recruitment of immune cells to the injury site. Caudal fin amputation was performed in the *Tg(coro1a:EGFP)* line in 3 dpf larvae, in which leukocytes are marked by EGFP expression. Larvae were treated with either 10 µM Sulfasalazine, 10 µM Etodolac, or 0.01% DMSO (used as a drug diluent control) for 5 hours (Figure S8A). Etodolac, but not Sulfasalazine, showed a significant reduction in immune cell recruitment at the injury front (p.val <0.0001). While Etodolac has a fast nonsteroidal anti-inflammatory action (Brocks & Jamali, 1994), Sulfasalazine requires its metabolization in the gut, taking a longer time to enter circulation (Choi et al., 2026), possibly explaining the difference in the observed “short-term readout” (Figure S8B,C).

To test whether alleviating inflammation in the paternal generation after cardiac damage could recover the phenotype in the next generation, we next injected 10 mg/kg Etodolac in adult cryoinjured males every 24 hours for 7 days. At 7 dpi, we crossed treated males with control females and assessed cardiac development in the next generation. As controls, we used cryoinjured males injected daily with only the solvent, 1% DMSO, and crossed with untreated females 7 days after (Figure 5G). The offspring of injured fathers treated with Etodolac showed an approximately 5% reduction in cardiac dimension with respect to the control (Figure 5H, I, Figure S9A,B). At the functional level, Etodolac did not recover fractional shortening (Figure 5J), but showed a statistically significant reduction in heart frequency and cardiac output (Figure 5K-L). Other cardiac parameters did not show alterations between the two groups (Figure S8C-E), but body length was significantly reduced in the offspring of injured fathers treated with Etodolac (Figure S8F), suggesting a generally slower development. Finally, we generated a heatmap with values normalized to the uninjured control group that allowed us to directly compare the effect of pro- and anti-inflammatory treatments on cardiac parameters with those arising from parental injury. In this map, we used cardiac measurements normalized to body size, therefore ruling out that the cardiac effect of the treatments is soley due to overall growth. This representation revealed that pro-inflammatory activation (*i.e.* HS of the IFNɣ transgenic line in the absence of injury) led to the development of a similar cardiac phenotype to that observed in the F1 of injured fathers. In contrast, anti-inflammatory treatment (i.e. Etodolac treatment after parental CI) induced the opposite effect, mirroring values obtained by the control groups (F1 of uninjured fathers, no HS treatment of the IFNɣ transgenic line, or HS of a non-transgenic parental line) (Figure 6M)

**Fig. 6:**
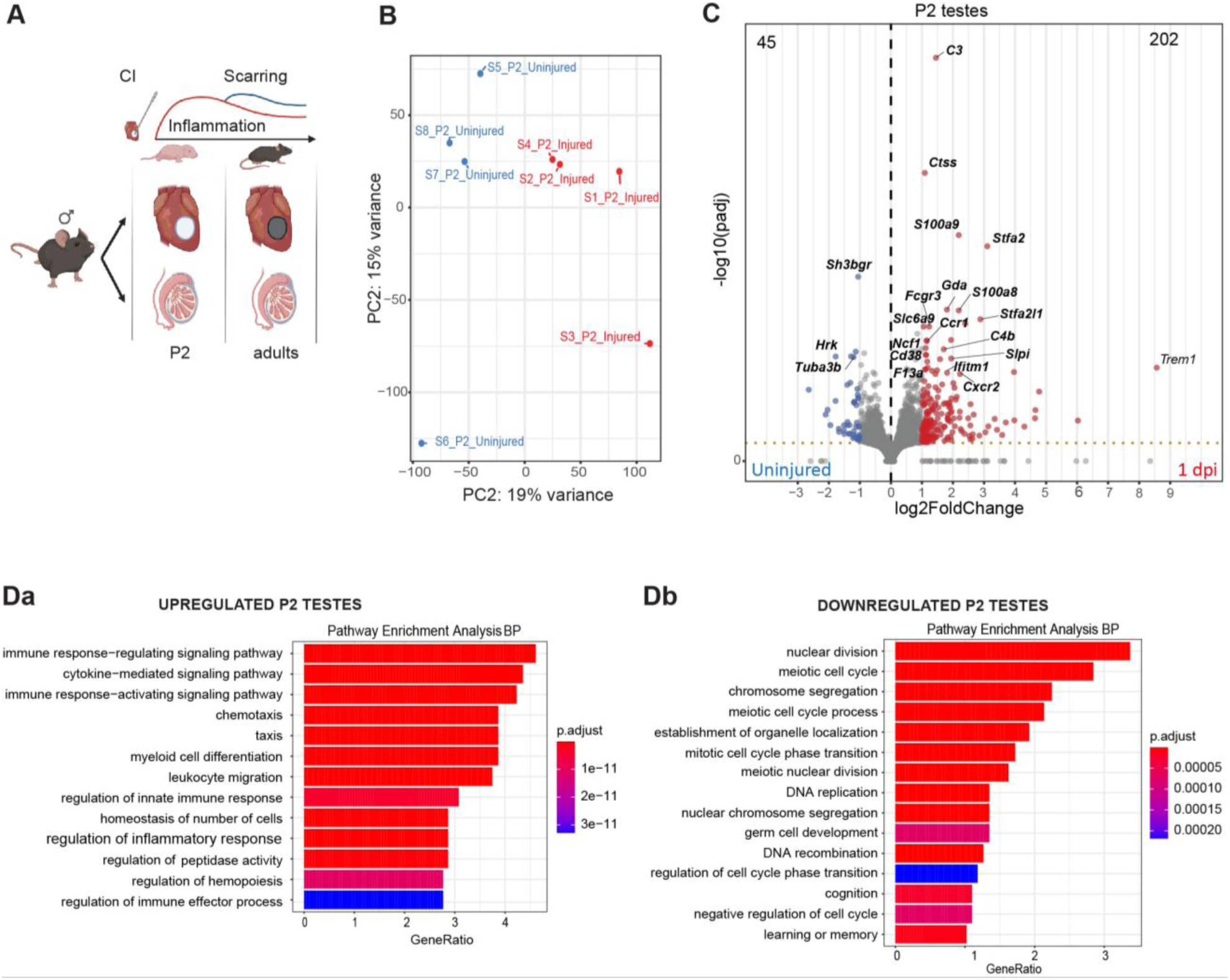
Neonatal cardiac CI leads to transient gene expression changes in mouse testes. (**A)** Experimental workflow: cardiac cryoinjury (CI) was performed at neonatal day 1 (P1). Testes were extracted from cryoinjured and uninjured mice one day after injury (P2), or once the animals reached adulthood. Bulk RNA-seq was performed on the collected samples. Image created in BioRender. Coppe, B. (2026) https://BioRender.com/6m9279f (**B)** Principal Component (PC) analysis of P2 samples. Each dot represents one biological replicate. blue, samples from uninjured animals; red, samples from injured animals. (**C)** Volcano plot of P2 testis from injured versus uninjured mice. Thresholds (|Log2FoldChange| ≥ 1, - log10(padj ≤ 0.05)). (**D)** Gene ontology pathway analysis of up (Da) and downregulated (Db) genes in testes collected from injured (ventricular cryoinjury at P1) *vs* uninjured animals at P2. RNA-seq results from adult testes are shown in Figure S10.

Together, these data suggest that sterile inflammation, which can be induced by cardiac injury or systemically activating IFNɣ in the paternal generation, influences cardiac development in the F1 generation, and anti-inflammatory treatment can partially recover this transmission (Figure 5M).

### A cardiac injury also elicits an early immune response in mouse gonads

Similarly to 3 dpf larvae born from injured zebrafish males, mice born from fathers who experienced neonatal CI early in life (*i.e.,* 1 day postpartum (dpp) = P1) exhibit cardiac enlargement under homeostatic conditions (*i.e.,* 3 weeks *postpartum* (wpp)) (Coppe et al., 2025).

To investigate whether a similar transmission mechanism is shared between these two species, we compared changes in gene expression in testes of uninjured and injured male mice. A CI was performed at P1. To distinguish acute cardiac injury responses from long-term adaptations associated with cardiac remodeling, we collected testes at one day after the injury (P2) and around eight months post-partum (adulthood) (Figure 6A). At P2, injured and uninjured samples segregated from each other in the PCA (Figure 6B); however, no changes were observed in testes collected from adults between the two groups (Figure S10A,B). We detected 274 DEGs at P2 (padj ≤ 0.05 and |Log2FoldChange| ≥ 1), of which 202 were upregulated in injured P2 males, and 45 were downregulated. Among upregulated genes, many were associated with the inflammatory response, including *Complement Component 3 (C3), Cathepsin S (Ctss), Fc Gamma Receptor IIIa (Fcgr3), Neutrophil Cytosolic Factor 1(Ncf1), C-C Motif Chemokine Receptor 1 (Ccr1), Complement C4B (C4b), Cd38, Secretory Leukocyte Peptidase Inhibitor (Slpi), Interferon Induced Transmembrane Protein 1 (Ifitm1),* and C-X-C motif chemokine receptor 2 (*Cxcr2*) (Figure 6C and <u>Bulk_RNA-seq_testes_M</u>). Among the top downregulated genes emerged *SH3 Domain Binding Glutamate Rich Protein* (*Sh3bgr)*, a gene specifically expressed in striated muscle and detected in mouse genitourinary tissues (PubChem, 2006.; Scartezzini et al., 1997), *Harakiri, BCL2 Interacting Protein* (*Hrk)*, promoting apoptosis (King et al., 2026), and *Tubulin Alpha 3B* (*Tuba3b)*, whose downregulation is associated with impaired germline cell maintenance and spermatogenesis (Fan et al., 2025) (Figure 6C).

GO biological pathways enrichment analysis performed on upregulated genes indicated the activation of the immune response (*i.e.* “immune response-regulating signaling pathway”, “cytokine−mediated signaling pathway”, “immune response−activating signaling pathway”, “myeloid cell differentiation”, “leukocyte migration”, *etc*.) (Figure 6Da), while GO biological pathways performed on downregulated genes found pathways presumably associated with the control of spermatogenesis (*i.e.* “nuclear division”, “meiotic cell cycle”, “chromosome segregation”, *etc*.) (Figure 6Db). GSEA enrichment analysis further confirmed the activation of inflammation-related pathways, and identified pathways associated with muscle development, suggesting a possible intercommunication between the damaged heart and the testis. Downregulated pathways were associated with the control of cell cycle and spermatogenesis (Figure S10C, <u>Bulk_RNA-seq_testes_M</u>).

Together, these data suggest that, similarly to what we previously observed in fish, cardiac injury transiently activates inflammatory responses and regulates spermatogenesis progression in the testis.

The transient alterations in gene expression observed in the neonatal mouse resembled the effect observed in the zebrafish, where 133 dpi gene expression changes in gonad also dissipated (Figure 1A, S1Bb). A transient change in gene expression could, however, lead to permanent changes at the level of chromatin. Since male mice reach sexual maturity and are typically bred from approximately 3 months postnatal onwards, we further investigated chromatin regulation in adult males’ gametes. Firstly, we analyzed changes in DNA methylation patterns in sperm. We used the Infinium Mouse Methylation BeadChip (Illumina), which allowed us to investigate 285K methylation sites per sample at single-nucleotide resolution. Similarly to the zebrafish, no major changes were observed at the DNA methylation level in the mature sperm collected from adult mouse males that underwent neonatal cardiac CI compared to uninjured males (Figure S11A). In fact, only 34 out of 296’070 (∼0.01%) probes were found differentially methylated (P.val ≤ 0.05; Effect size: 10%) between the two conditions (Figure S11B,C, DNA_Met_sperm_M), while no DMR was discovered.

We next investigated chromatin accessibility changes using single nuclei (sn) ATAC-seq of whole testis (Figure 7A). 64’335 nuclei were captured from seven samples, *i.e.* testes from 4 controls and 3 injured (P1) mice, and sequenced (Figure S12A). We obtained a median of accessible genes per nucleus ranging from 2’419 to 10’655 (Figure S12B). Eleven main cell clusters were identified (Figure 7B). Clusters and subclusters were assigned to specific cell populations using established gene markers associated with chromatin accessibility (Kavarthapu et al., 2023; Suen et al., 2023; Tan et al., 2020; Whelan et al., 2025; Y. Yin et al., 2015) (Figure 7C, see materials and methods). The proportion of cells was similar between conditions across clusters (Figure S12C), suggesting no permanent alterations in cell populations are present in adults following neontal cardiac injury.

**Fig. 7.**
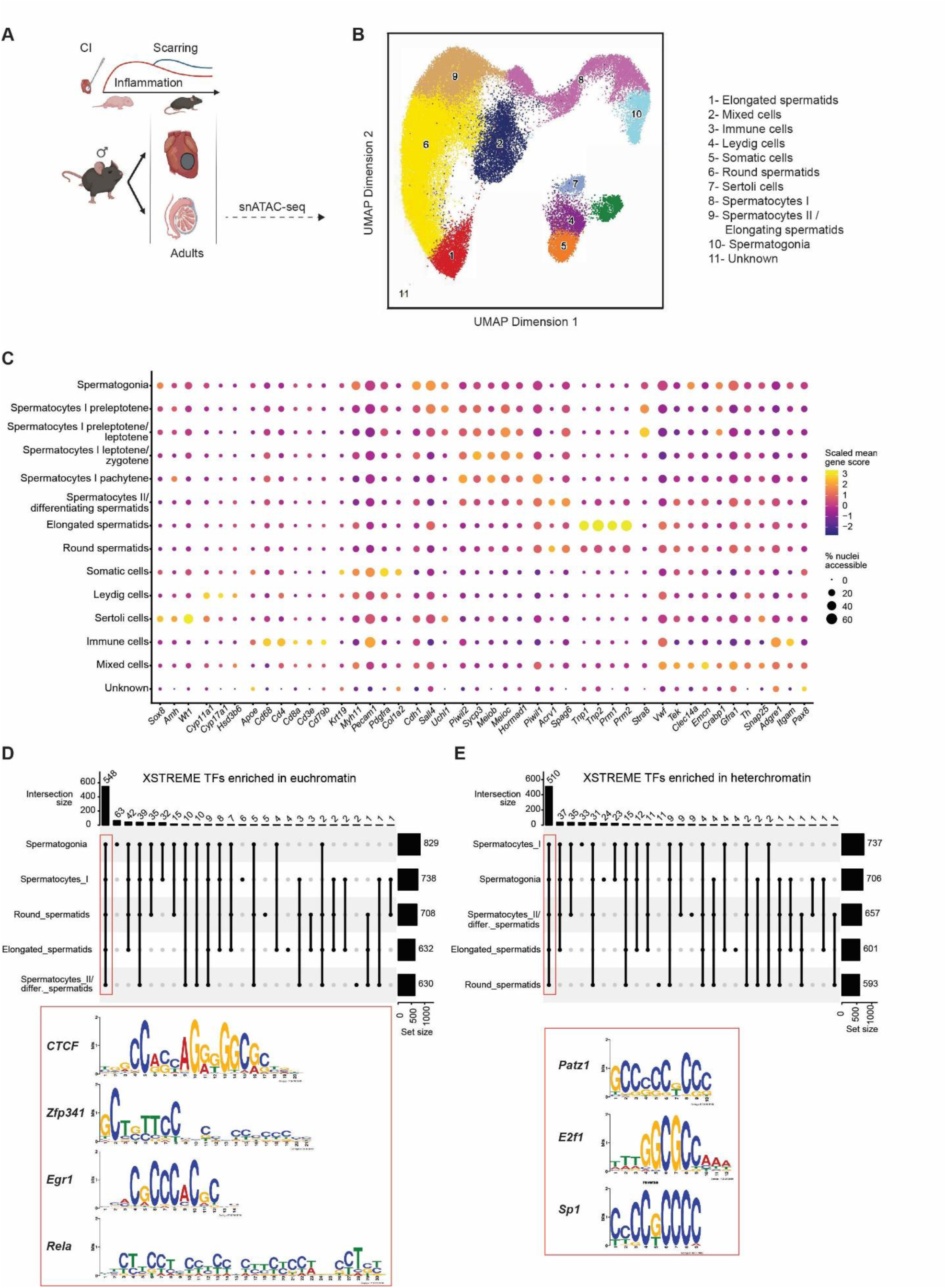
Single-nuclei chromatin accessibility assay in adult murine male gonads following neonatal heart damage. (**A)** Schematic representation of sample collection. Adult mouse testes were collected from animals that underwent neonatal cryoinjury (CI) and from control uninjured animals. Samples were processed for scATAC-seq. Illustration created in BioRender. Coppe, B. (2026) https://BioRender.com/6m9279f (**B**) Uniform Manifold Approximation and Projection UMAP displaying identified main clusters of male gonads’ cell populations. (**C**) List of markers used for cell type clustering. (**D)** Up**s**et graphs reveal shared putative TFs among different cell types differentially enriched in euchromatic (Da) and heterochromatic (Db) regions. Binding sites for Patz1, E2f1, Sp1 were enriched in heterochromatin regions, and CTCF, Zfp341, Egr1, Rela within euchromatin regions, among others.

We focused our attention on the analysis of differentially accessible regions (DARs; P.val≤0.05, |logFoldChange| ≥ 0.585) in germ cells between conditions and performed pathway enrichment analysis. Spermatogonia displayed the highest number of DARs (2’958), possibly due to the highly accessible chromatin state of this cell type associated with stem cell maintenance and differentiation potential (<u>snATAC-seq_testes_M</u>). We also identified DARs in spermatocytes Type I and II as well as in spermatid stages. The presence of DARs in spermatogonia suggests stable alterations maintained throughout life following neonatal cardiac damage, whereas DARs in the elongated spermatids might contain information likely transmitted to the sperm and fertilized egg. We were not able to identify DARs maintained in the same *locus* throughout spermatogenesis (Figure S12D-G), likely reflecting the highly dynamic chromatin remodeling occurring during this process. This might also be a consequence of the clonal dynamics observed during spermatogenesis that preclude precise clustering of a full spermatogenesis cycle at the clonal level with the current approach. We performed pathway analysis of genes associated with DARs found in either heterochromatic or euchromatic states in the testis of injured *versus* uninjured males for all main germ cell clusters (Figure S13A-H). Gene *loci* related to biological processes involved in sperm maturation, such as cilium organization, were found in heterochromatin, and therefore presumably transcriptionally silent, in most germ cell populations of injured animals (*i.e.* spermatocytes I, round, and elongated spermatids) (Figure S12C, F and H). In round spermatids, gene *loci* involved in the negative regulation of TOR signaling were also found in a heterochromatic state (Figure S12F), indicating potentially dysregulated spermiogenesis (Sahin et al., 2018). Chromatin regions associated with *Wnt* pathway genes were found enriched in euchromatic regions in both spermatogonia and spermatocytes I of injured males (Figure S12B,D), suggesting promotion of division and differentiation into specialized sperm cells rather than maintenance of the stem cell pools in this group (Takase & Nusse, 2016). The results are in agreement with proteomics analysis performed in zebrafish, where we also observed dysregulation of pathways related to cell division (Figure 1G,H). Together, these data reflect the maintenance of a “structural memory” of gonadal response occurring early-life in repsonse to neonatal cardiac damage.

We next performed enrichment analysis of TFBS (P.val<0.05) using Xstreme from Meme Suite software. Remarkably, the majority of putative TFs binding DARs in both heterochromatin (LogFoldChange<0) and euchromatin (LogFoldChange>0) states in samples from injured animals were shared between all germ cell clusters, suggesting that common transcriptional regulators are maintained across developmental stages (Figure 7D-E), even though the underlying accessible chromatin regions differed (Figure S12D-G). TFs enrichment analysis identified TFs involved in stemness (HES family, HEY1/2) and cell cycle control (TFs part of E2F family), epigenetic/chromatin regulation (CTCF, CTCFL, DNMT1, TET1, KMT2A, KDM2A/B, PRDM family among others), development (HOX family), immune response (Rela/b, NFKB1/2, IRF and SMAD family) and hypoxia (HIF1a) (<u>snATAC-seq_testes_M</u>). Among putative TFs enriched in injured testis (*i.e*. euchromatin), and shared across all spermatogenesis states were *CTCF*, *Zfp341 (Zinc Finger Protein 341*), a TF critical for immune system regulation controlling *STAT3* activation (Frey-Jakobs et al., 2018), *Egr1 (Early Growth Response 1)*, induced by growth factors, inflammation, and cellular stress (Lim et al., 1998), and *Rela*, involved in immune response and inflammation (Ouaaz et al., 1999) (Figure 7D). Among the top putative TFs enriched in uninjured samples (*i.e*. heterochromatin), we found multiple TFs involved in the regulation of spermatogonia maintenance/proliferation and spermatogenesis regulation, such as *PATZ1* (*POZ/BTB And AT Hook Containing*) (Fedele et al., 2008), *E2f1* (*E2F transcription factor 1*) (Hoja et al., 2004), and *Sp1* (*Sp1 Transcription Factor*) (Persengiev et al., 1996) (Figure 7E, <u>snATAC-seq_testes_M</u>). Together, molecular analyses performed in mice suggest that cardiac injury induces an inflammatory response in male gametes 24 hours post-injury (Figure 6C,D), leading to long-lasting changes in chromatin features related to spermatogenesis, chromatin regulation, and inflammation (Figure 7).

### Transcriptomic gene signature of an inflammation memory in F1 mice hearts from injured fathers

In F1 adult hearts from injured zebrafish males, we have observed alterations in gene expression associated with inflammation (Figure 3B,C). We wanted to address whether gene expression was also affected in hearts collected from F1 mice from fathers that underwent cardiac CI. We therefore performed bulk RNA-seq of the whole heart at 3 wpp (Figure 8A). Samples were collected from males (four controls and three experimental) and females (two controls and three experimental) obtained from three different crosses per condition. Following PCA analysis, we excluded sample “Injured P0_16” statistically categorized as an outlier (Figures S14A-B). Differential expression analysis identified 24 DEGs (padj≤0.05 & |Log2FoldChange| ≥ 0.58), of which 15 were upregulated and 9 downregulated in the heart of offspring from injured fathers under physiological conditions (Figure 8B-C, <u>Bulk-RNA-seq_F1_heart_M</u>). Among the upregulated genes were *Tnf (Tumor Necrosis Factor)*, *Il31ra (Interleukin-31 receptor A)*, and *Ccr5 (C-C motif chemokine receptor 5),* previously shown to be involved in inflammatory processes and cardiac hypertrophy (Wei et al., 2020; Liu et al., 2023; Kroll-Palhares et al., 2008; Dobaczewski et al., 2010; Bristow, 1998). The downregulated genes were *Ciart (Circadian Associated Repressor Of Transcription,* also known as *Chrono)* and *Per2 (Period Circadian Regulator 2)*, two circadian-associated genes (Anafi et al., 2014; M. Kim et al., 2018) that may be implicated in maladaptive cardiac function (Lin et al., 2023), *Tnnt3 (Troponin T3, Fast Skeletal Type)*, whose variant causes contraction deficiencies and dilated cardiomyopathy (Jadidi et al., 2025), and *Trp53inp1 (Transformation-related protein 53-inducible nuclear protein 1)*, a highly conserved stress-response gene involved in the inhibition of cardiomyocyte mitosis (Cai et al., 2018), among others (Figure 8C).

**Fig. 8.**
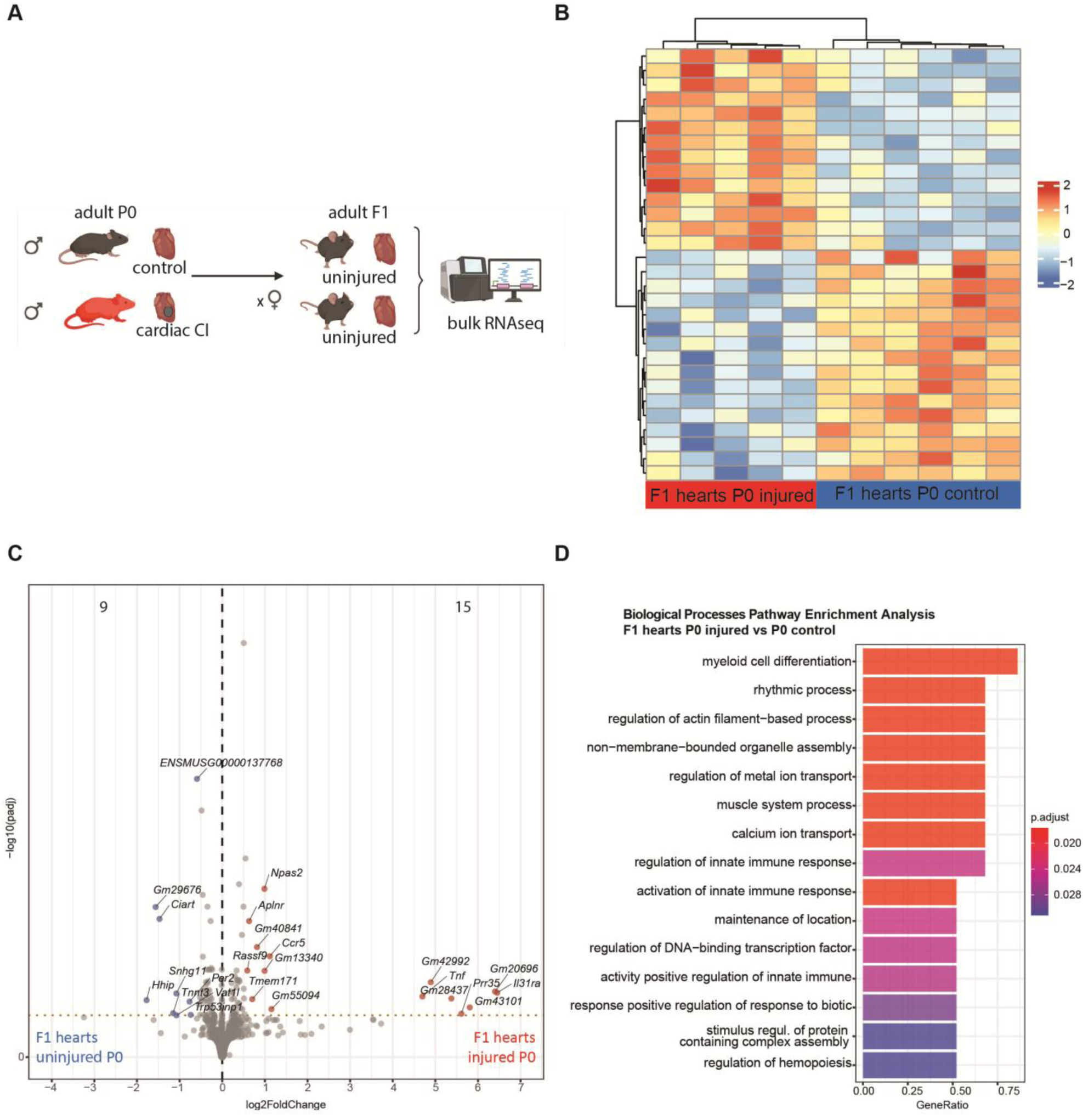
Effect of a paternal cardiac injury on gene expression in F1 hearts. **(A)** Schematic representation of experimental outline. Mice were injured at neonatal day 1 (P1). Once adult, males were crossed with uninjured females. For the control group, mice were grown in the same conditions from parallel crosses. Again, once adult, the males were crossed with the same colony of uninjured females. Hearts from 3-week-old F1 animals were collected for RNA extraction and sequencing. Image created in BioRender. Coppe, B. (2026) https://BioRender.com/6m9279f (**B**) Heat map of the 30 top differentially expressed genes between experimental (F1 from injured fathers) and control (F1 from uninjured fathers) groups. (**C**) Volcano plot. Red and blue dots mark differentially expressed up- and downregulated genes between F1 hearts from injured and uninjured males. Padjust ≤0.05. (**D**) Gene Ontology Biological Process Enrichment Analysis of RNA-seq data.

GO biological enrichment pathway analysis identified pathways associated with immune response such as myeloid cell differentiation and regulation/activation of innate immune response, and pathways related to cardiac contractility, including rhythmic process, regulation of actin filament-based process and regulation of metal/calcium ion transport (Figure 8D, Bulk-RNA-seq_F1_heart_M).

Altogether, transcriptomic data of mouse hearts collected at 3 wpp from the offspring of injured fathers under homeostatic conditions suggest a low-grade activation of the inflammatory response, similarly observed in the zebrafish, and a gene signature associated with contractile alteration, possibly associated with functional alterations observed by echocardiography at this time point (Coppe et al., 2025).

## DISCUSSION

Growing evidence indicates that consequences of the interaction between the environment and an exposed individual do not necessarily terminate when the triggering stimulus ends but can extend across one or multiple generations (Skvortsova et al., 2018; Fitz-James & Cavalli, 2022). Not only exposure to external *stimuli*, such as chemicals, altered diets, or traumatic experiences, but also to internal alterations, such as organ damage (Zeybel et al., 2012) and sepsis (Bomans et al., 2018) seem to have the potential to influence next generations’ phenotypes.

Parental CVDs increase offspring’s risk of developing the same disease, and multiple factors, such as genetic predisposition and lifestyle, have been suggested to contribute to this inheritance (Hossin et al., 2024). Here, zebrafish and mice were used as research models to understand the intergenerational influence of paternal cardiac damage alone. In the event of heart failure, inter-organ crosstalk between the heart and peripheral organs has been reported both in zebrafish (Sun et al., 2022) and humans (Rettkowski et al., 2025b; Schiattarella & Kontaridis, 2025) and systemic inflammatory responses across multiple organs have been observed after MI in both zebrafish and mice (Cortada et al., 2024). Similarly, following cardiac damage, we have observed inflammation in the male gonads of both species. Our transcriptomics analysis identified temporary gene activation in response to cardiac damage in testes, followed by recovery to a normal transcriptional state, regardless of whether the heart regenerated (in zebrafish) or developed permanent scarring (in mice). Alterations in the dynamics controlling spermatogenesis were observed at the molecular level in both species. Interestingly, many pathways found associated with inflammation, such as *mTOR*, *JAK/STAT*, and *NF-κB*, are also involved in cell proliferation and could ultimately play a dual role in spermatogenesis progression.

Changes in DNA methylome and chromatin accessibility were assessed in the germ line of both species to understand whether the altered local environment would influence chromatin regulation of male gametes, possibly leading to the transmission of newly established gene-regulatory states. In zebrafish, after fertilization, maternal DNA methylation matches paternal patterns during epigenetic reprogramming, making this epigenetic marker a likely candidate for *patriline* inheritance (Potok et al., 2013; Wu et al., 2011). In mice, DNA methylation can be transmitted from parents to their offspring (Takahashi et al., 2023). While no DNA methylation regions were found to be altered in sperm after cardiac damage, neither in zebrafish nor mice, chromatin accessibility changed and reflected the gonad’s inflammatory state and altered control of spermatogenesis in both species.

In mouse sperm, where ∼95% of histones are replaced by protamines (Luense et al., 2016), even a small amount of contaminating DNA from other cell sources can contribute to a major fraction of the nucleosomal signal and potentially overcome sperm-derived chromatin signals (Q. Yin et al., 2023). In zebrafish, by contrast, germ cells retain histones throughout spermiogenesis, so that external minor chromatin contaminations would not majorly affect sperm nucleosomal signals. For these reasons, to assess chromatin alterations established following cardiac injury and possibly transmitted to the next generation, we used bulk ATAC-seq in zebrafish sperm, whereas in mouse, we relied on single-cell ATAC-seq of testis. In zebrafish, both binding sites for *Rela*, the rapid canonical *p50/RelA* pathway of *NF-kB*, responsible for immune response, inflammation, and apoptosis regulation (Mao et al., 2025), and *Nr2c2*, a nuclear receptor essential for normal spermatogenesis (Mu et al., 2004), were found enriched in chromatin accessible regions in the sperm following cardiac injury. *CTCF*, a master regulator of chromatin architecture, was also enriched, pointing to a broader restructuring of sperm chromatin architecture after cardiac injury. Several zinc finger TFs showed enrichment across both active and silenced chromatin regions, which may reflect changes in zinc availability in testes, associated with increased oxidative stress and inflammatory signaling (Prasad, 2008). Similarly, in mice, we identified accessible regions enriched for *CTCF*, Rela, and *PATZ1* binding sites, among others, suggesting conserved chromatin remodeling, inflammatory, and spermatogenesis-related responses to cardiac injury across species. Interestingly, CTCF has been already shown to be responsible for the regulation of transgenerational inheritance mechanisms in mice (Jung et al., 2022). Here we identified the presence of motifs of the same TF in DARs throughout spermatogenesis and its conserved presence in germ cells between two species. However, we did not observe a specific permanently imprinted locus in differentiating germ cells in response to a cardiac injury. This might be the result of the lack of true clonal germ cell analysis or can also mean that there is not a single locus involved in transmission of information, but rather an overall chromatin rearrangement with consequences for the next generation.

Chromatin accessibility is regulated by the binding of transcription factors, chromatin remodelers, and architectural proteins, as well as by DNA methylation and histone PTMs. Previous studies have shown that alterations in histone PTMs in gametes can also correlate with phenotypic changes in the next generation, including in zebrafish (González-Rojo et al., 2019; Lombó et al., 2019). Thus, future analysis focused on the transmission of histone PTMs through sperm and early embryos could further elucidate the mechanisms underlying the intergenerational effects of paternal cardiac injury.

Previously, different models of paternal inflammation have been reported to influence offspring growth, regeneration (Z. Zhang et al., 2020), immune response (Bomans et al., 2018), and metabolism (Y. Zhang et al., 2021). We investigated whether paternal cardiac damage-induced inflammation influences the cardiovascular system in the next generation under homeostatic conditions. In zebrafish, cardiac overexpression of *Isg15* and *Rsad2* suggested dysregulation of interferon signalling pathways, whereas in mice, key inflammatory genes, including *Tnf* and interleukin/chemokine receptors, similarly indicated immune activation. Taken together, these results reveal a basal inflammatory signature in the heart of offspring from injured fathers in both zebrafish and mice. Early in life (3 dpf and 3 wpp, in zebrafish and mice, respectively), both species showed features associated with ventricular dilation under homeostatic conditions. Interestingly, similar cardiac phenotypic alterations were observed, among other features, in the offspring of zebrafish males exposed to Bisphenol A (Lombó et al., 2015). Future analyses of cardiac morphology and function following injury in the F1 generation may help determine whether the observed low-grade basal inflammation influences cardiac damage responses in these two species.

In zebrafish, we confirmed the role of inflammation in the intergenerational transmission of paternal cardiac damage, using two approaches: simulating an inflammatory state or treating the paternal generation with anti-inflammatory drugs and subsequently assessing cardiac phenotype in the F1 generation. We did not extend anti-inflammatory treatment to mice, since post-infarct administration of systemic anti-inflammatory drugs has been associated with impaired cardiac repair and adverse outcomes (*i.e*. adverse remodeling, arrhythmogenesis, ventricular rupture), raising our concern about administering anti-inflammatory treatments to mice following myocardial infarction (Huang & Frangogiannis, 2018). However, it is likely that other severe surgical interventions that trigger systemic acute inflammation, such as thoracotomy (Talbot et al., 2015), may also influence the germ line and contribute to cardiac phenotypic alterations in the next generation. Notably, F1 offspring of sham-operated male mice have been reported to exhibit cardiac phenotypes similar to those observed in offspring of injured males, further supporting a role for paternal inflammation in intergenerational inheritance in mammals (Coppe et al., 2024). Similarly, we estimate that other severe tissue injuries to the parental generation could also elicit alterations in cardiac function in offspring. While this study focused on the intergenerational effects of paternal cardiac damage on the next generation’s heart, other organs are also likely to be affected.

Through this study, we showed that cardiovascular disease transmission is influenced not only by lifestyle or genetic predisposition but also by cardiac damage itself. With more men having children later in life and myocardial infarction increasingly affecting younger adults (Hossin et al., 2024), our study highlights the importance of further exploring the molecular mechanisms underlying the transmission of inflammatory diseases and cardiac dysfunction.

## MATERIALS AND METHODS

### Zebrafish husbandry

All experiments were performed using zebrafish (Danio rerio) embryos, larvae, and adults (4–17 months old) at the Institute of Anatomy of the University of Bern. Adult fish were maintained at a maximum density of 5 fish/L under standardized conditions: 27.5–28 °C, 14:10 h light:dark cycle, conductivity of 650–700 μS/cm, pH 7.5, and 10% daily water exchange. Fish were fed two times daily with one feeding of Artemia (Ocean Nutrition) and one feeding of dry food (ZM-000, Gemma Micron 150, and 300 for larval, juvenile, and adult stages, respectively). All animal procedures were approved by the Animal Care and Experimentation Committee of the Canton of Bern, Switzerland (licenses BE46/19, BE109/2021, BE05/2025), and were conducted in accordance with Swiss regulations. Zebrafish strains used in this study included wild-type AB, *Tg(myl7:EGFP)* (González-Rosa et al., 2011b), *Tg(myl7:mRFP)*^ko08^ (Rohr et al., 2008), *Tg(hsp70l:ifng1-2-V5,cryaa:Cerulean)*^s994T^ (Sawamiphak et al., 2014), and the Tg(*coro1a:eGFP*)^hkz04tg^ (L. Li et al., 2012). In the paternal generation, replicates for both experimental and control groups were established by pooling male siblings, with each replicate comprising individuals derived from different parental crosses. For intergenerational experiments (F1 generation), males from the paternal generation (siblings) were separated at 5 dpf into different tanks, with injured and uninjured groups maintained separately. Females of similar age, also consisting of sibling groups, were selected for the maternal generation. Consequently, within each replicate, F1 individuals were full siblings, whereas individuals across replicates and conditions were 3^rd^ degree relatives.

### Mouse husbandry

Experiments conducted in mice were performed in neonatal and up to one year old C57BL/6J mice. Animals were raised together until weaning (3 wpp); afterwards, they were maintained in a density of 4 animals/cage, except bred males, which were isolated after the first breeding. All experiments were performed in a Specific Pathogen Free facility under the following conditions: 20-24°C, 12 hours light-dark cycle, and 45-65% relative humidity. Standard food and water were available *ad libitum* and, together with the litter, were changed regularly. All experiments were performed in the Spanish National Centre for Cardiovascular Research (CNIC) in Madrid (Spain) and approved by the Community of Madrid “Dirección General de Medio Ambiente” in Spain. All animal procedures are conformed to EU Directive 86/609/EEC and Recommendation 2007/526/EC regarding the protection of animals used for experimental and other scientific purposes, enforced in Spanish law under Real Decreto 1201/2005. Experiments were conducted under the license PROEX 310_19.

The parental generation was formed by injured (CI at P1) or uninjured males that were bred at adult age with control sibling females of similar age. Uninjured and injured males were not siblings among them, since mothers would eat unhealthy-looking mice if pooled together with uninjured animals. The F1 generation was composed of siblings from different families.

### Zebrafish ventricular cryoinjury

CI of the ventricle was performed in adult fish as described by (Marques et al., 2021). Briefly, fish were anesthetized in 0.032% Tricaine (wt/vol) and put belly up inside a foam. The surgery was performed under a dissecting microscope by incising skin, pectoral muscles, and pericardium with a single small cut performed with a microdissection scissor. The ventricle was exposed by gently squeezing the abdomen and dried with a piece of tissue paper. A copper probe (0.5 mm diameter) previously frozen in liquid nitrogen was placed on top of the ventricle apex for three seconds and removed. The belly compression was terminated to allow the heart to go back to its original position, the juxtaposing skin was pulled back to help close the injury, and the fish was put back in fresh water. Animals were finally woken up by gently pipetting fresh water on the gills for a few seconds.

### Mouse neonatal left ventricular cryoinjury

Before the surgery, a few drops of surgical glue were spread inside the cage, close to the net, to desensitize the mother to its smell. Afterwards, half of the litter was separated by the mother, and the surgery was performed. After waking up and recovering body temperature, the pups were put back with the mother, and the other half of the litter was processed to limit mother stress caused by pups’ separation. The surgery was performed as described by (Zhao et al., 2021). Briefly, P1 mice were anesthetized by induced hypothermia and maintained in a supine position on top of a Petri dish filled with ice. A small incision was made in the skin over the approximate projection of the 3rd and 4th intercostal space. Then the pectoral musculature was dissected, exposing the thoracic cage. An additional incision was performed at the level of the 4th intercostal space to expose the interior of the rib cage using fine curved blunt forceps. By applying slight pressure to the abdomen, the apical region of the heart was exposed. A 24-gauge (1 mm) probe, previously immersed in liquid nitrogen, was then applied to the heart, freezing approximately 15% of the cardiac tissue. The procedure was completed by closing the costal margin with an 8-0 silk suture, followed by the administration of a 5 μL drop of buprenorphine (0.03 mg/mL) into the subcutaneous pocket. The skin incision was then sealed using surgical glue. Mice were placed on a thermostatically controlled heating plate at 38°C for 10 minutes to restore body temperature and, once normal breathing had resumed, were returned to their mother.

### Zebrafish and mice euthanasia

Zebrafish were euthanized by an overdose of Tricaine (0.16% w/v). Mice were euthanized by decapitation when neonatal (P2) or by CO₂ overdose when aged 3 wpp or older.

### Zebrafish sperm collection

Selection of fertile fish was performed by breeding adult males before sperm collection. Sperm was collected in the morning in uninjured and 7 dpi males in anaesthetized fish (0.016% Tricaine), with the help of a 10μl capillary and aspirator tube. Shortly, males were anaesthetized in 0.032% Tricaine (wt/vol) and placed ventral side up inside a sponge. The anal fins were opened to expose the urogenital papilla, which was then gently dried with tissue paper. The capillary was then placed on top of the cloaca, and the ejaculate was gently collected by applying some pressure close to the urogenital papilla. The capillary was finally removed, 0.3-0.5 μL of sperm was resuspended in a solution containing 0.5% BSA (wt/vol), 5mM CaCl_2_, and 5mM MgCl_2_ in PBS on ice, and the animal was awakened in fresh water. Sperm from five siblings were pulled for each sample. Samples were then centrifuged for 10’ 1500rcf at 4°C and supernatant discarded. Pellet was resuspended in PBS and washed twice prior to use. Purity of sperm was examined under the microscope.

### Zebrafish testes collection, Hematoxylin & Eosin staining, and analysis

Testes were dissected from adult zebrafish following tricaine overdose. Samples were fixed overnight at 4°C in 2% paraformaldehyde (PFA). For paraffin embedding, testes were included per block after dehydration through a graded ethanol series (70%, 90%, 2×100%) and xylene (2×100%) and embedded in paraffin. Paraffin blocks were sectioned at 7 μm using a microtome to obtain representative sections of the whole testes. Paraffin sections were deparaffinized, rehydrated through graded ethanol, and washed in distilled water. Sections were stained with Haematoxylin for 5 min and slides were rinsed with water for 5 min prior to 1 min Eosin staining. Slides were rinsed again with distilled water, dehydrated through graded alcohols, and cleared in xylene prior to mounting. Images were acquired using Hamamatsu NanoZoomer 2.0RS slide scanner and processed with NDP.view2 image viewing software (Hamamatsu). At least two seminiferous tubules in a minimum of two sections were analysed per animal. Within each seminiferous tubule, spermatocysts (i.e., spermatogonia, spermatocytes, and spermatids) were identified. The average number of each spermatocyst type was calculated per animal and expressed as a percentage of the total spermatocysts.

### 1K-cells zebrafish embryos collection and processing

Wild-type male fish were crossed with wild-type control females before the injury or 7 dpi. Eggs deposition was verified and limited to 20 minutes. 3 hours post fertilization, 1K-cells embryos were collected and staged under the microscope. Dechorionation was performed by treating embryos with Pronase 2 mg/mL in E3 1 for 4-5 min at RT. Once the chorion was excluded, embryos were washed twice in E3 medium and collected in an Eppendorf tube. Water was eliminated, and samples were snap frozen and stored at -80°C until use.

### Mice testes and sperm collection

Sexually mature male mice were euthanized by CO₂ overdose. Ethanol was applied to the abdomen and hind limbs to prevent loose fur from adhering to surgical instruments and contaminating the dissection material. Using microsurgical scissors, the skin and abdominal wall were incised to expose the reproductive organs. Testes and epididymides were collected using forceps and separated in a sterile Petri dish containing PBS. Testes were snap-frozen and stored for RNA extraction or further processed for snATAC-seq.

Sperm was isolated from the cauda epididymides as previously described by Jung et al. (2022). Briefly, both cauda epididymides were cleaned by removing surrounding fat and blood vessels, transferred to a 30-mm Petri dish, covered with a drop of Donners medium (Hisano et al., 2013), and minced using a syringe needle. The tissue was immediately transferred to a round-bottom tube containing 8.5 mL of Donners medium and incubated at 37°C for 1 h to allow spermatozoa to swim out of the epididymal tissue. The upper fraction, containing motile spermatozoa, was then collected and passed through a 40-µm cell strainer. Sperm purity was assessed by light microscopy.

### Zebrafish echocardiography

Prior to echocardiographic assessment, animals were assigned to tanks labelled with colour-coded identifiers to ensure blinding. The operator performing the echocardiography was unaware of group assignments. Unblinding was conducted during the data plotting stage.

Echocardiography was performed as described in (González-Rosa et al., 2014) with a few modifications. The day of the echocardiography, animals were anesthetized using a combined solution of 60 µM tricaine and 3 mM isoflurane dissolved in fish tank water, which prolongs anaesthesia duration while minimizing cardiac rhythm side effects.

After confirming full anesthesia (∼3 min), fish were immobilized ventral side up in a foam holder within a glass container filled with water and anesthetic maintained at 28°C to reduce stress in the fish.

The probe used for image acquisition was positioned ∼3 mm from the skin inside the water and covered with ultrasound gel to avoid interface artefacts. Transthoracic echocardiography was performed blinded by an expert operator using a high-frequency ultrasound system (Vevo 3100, FUJIFILM VisualSonics, Inc.) with a 70-MHz linear probe. Two-dimensional (2D) echography images were acquired at a frame rate above 320 frames/sec, and pulse wave Doppler (PW) was acquired with a pulse repetition frequency of 40 kHz. After measurement, fish were awakened in a recovery tank in fresh fish water.

Images were transferred and analyzed using the Vevo LAB 3.1.1 software (FUJIFILM VisualSonics, Toronto, Canada). Ventricular end-diastolic and end-systolic areas were manually measured from longitudinal-axis B-mode images. Heart rate was determined using pulsed-wave Doppler signal and averaged for 3 heartbeats.

Ventricular end-diastolic and end-systolic volumes, ejection fraction, fractional shortening, stroke volume, and cardiac output were automatically calculated using the “Area–Length” method offered by the Vevo LAB 3.1.1 software.

### High content imaging of larval hearts and data analysis

Uninjured and injured (7 dpi) wild-type AB males were paired-crossed simultaneously with transgenic females expressing a fluorescent protein under the cardiomyocyte (CM) promoter Myl7 (*i.e. Tg(myl7:EGFP)* (González-Rosa et al., 2011b), *Tg(myl7:mRFP)*^ko08^ (Rohr et al., 2008). In all experiments, both groups (F1 from uninjured and injured fathers) were crossed with mothers of the same transgenic background. In detail, paired adult fish (one male and one female) were placed inside a breeding tank and kept separated by a divider overnight. The divider was removed the following morning to allow spawning. Eggs were collected within 20 min after the removal of the divider and incubated at 28.5°C in E3 medium (5 mM NaCl, 0.17 mM KCl, 0.33 mM CaCl2, 0.33 mM MgSO4) with ∼0.001% methylene blue. After 24 hours, embryos were cleaned, and the medium was replaced with E3 medium containing 0.003% PTU to inhibit pigmentation. Embryos were selected based on the marker of interest (*i.e.* fluorescent heart) at 48 hpf. At 72 hpf, embryos were anesthetized with 0.2 mg/ml tricaine and positioned ventral side down in agarose beds in 96-well plates using 3D-printed templates, as previously described (Ernst et al., 2023).

Imaging was performed using the automated widefield fluorescent microscope Acquifer Imaging Machine (Bruker). An overview image of each well was acquired in brightfield using the 2X objective (pre-scan). Subsequently, the 10X objective was used to acquire a time series of 300 frames of the head region focused on the heart. Heart images were acquired using the FITC or TRITC fluorescence channels.

Body length and cardiovascular parameters were quantified in ImageJ and using the U-Net–based deep learning model HeartSeg (Ernst et al., 2023), which was updated to extract more cardiac parameters including Systolic and Diastolic cross-sectional area, Circularity, End Diastolic Volume (EDV), End Systolic Volume (ESV), Stroke Volume (SV), and Cardiac Output (CO). The updated version of HeartSeg allows visual inspection of segmentation. Only data correctly segmented for ventricle were kept for further statistical analysis.

### Caudal fin amputations in zebrafish larvae and anti-inflammatory drug treatment

Caudal fin amputation was performed on 3 dpf larvae of the Tg(*coro1a:eGFP*)*hkz05t* line (L. Li et al., 2012). Anesthetized larvae (0.2 mg/ml tricaine) were laid on a glass slide, and the caudal fin was amputated using a sterile scalpel. Immediately after amputation, larvae were maintained in petri dishes with the anti-inflammatory drugs Etodolac (10 µM), Sulfasalazine (10 µM), or 0.01% DMSO. Accumulation of *coro1a*:eGFP+ cells at the amputation site was recorded at 5 hours post-amputation (hpa) using 20X objective, and the FITC channel of the Aquifer machine (Bruker). Serial images spanning the entire fin thickness were acquired and processed as maximum intensity Z-projections using Fiji (ImageJ). Brightness, contrast, and color levels were adjusted for optimal visualization. Cell migration (number of *coro1a*:eGFP+ cells) was quantified within a 100 µm region adjacent to the amputation site, corresponding to the blastema.

### Anti-inflammatory drug treatments in adult zebrafish

Etodolac was dissolved in DMSO at a stock concentration of 25 mg/ml and used at a final concentration of 0.25 mg/ml. Injection volumes were adjusted according to fish weight to deliver 10 mg/kg intraperitoneally (IP). Etodolac was dissolved in DMSO at a stock concentration of 25 mg/ml and subsequently diluted to 0.25 mg/mL in 1% DMSO in PBS, with 10 mg/kg administered per fish. Fish were treated daily for 6 days, with injections performed every 24 hours in anesthetized males. The first injection was given immediately after cardiac injury. Seven days post-injury, males were paired with untreated females of the *Tg(myl7:EGFP)* (González-Rosa et al., 2011b), or *Tg(myl7:mRFP)*^ko08^ (Rohr et al., 2008) to generate F1 offspring. The same transgenic line was used for the control and experimental group within the same experimental replicate. For the control group, the same procedure was followed, except that 1% DMSO was injected instead of the anti-inflammatory drugs at the corresponding time points.

### Heat shock of adult zebrafish

Male fish from the *Tg(hsp70l:ifng1-2-V5,cryaa:Cerulean*)*s994Tg* (Sawamiphak et al., 2014) were subjected to a heat pulse by transferring them into a pre-warmed tank and maintaining them in an incubator at 38°C for 1 hour. Seven days after the heat shock, males were paired with untreated females of the *Tg(myl7:EGFP)* line to generate F1 offspring. Selected eGFP+ F1 larvae were imaged at 3 dpf using the Acquifer system to assess cardiac development, as previously described. In the control groups, the same procedure was followed using either fish lacking the heat-shock promoter (*i.e.* AB or *Tg(isg15:EGFP,myl7:EGFP*)) or fish from the same line that were not subjected to heat shock.

### Omics samples preparation, processing and bioinformatic analysis

Details of all omics experiments can be found in the supplementary materials and methods, including: RNA extraction and bulk RNA-seq of zebrafish and mice P0 testes and F1 hearts; mass spectrometry data processing and protein identification of zebrafish testes; DNA extraction and Whole Genome Bisulfite Sequencing of zebrafish sperm and 1k-cells embryos; DNA methylation analysis of mouse spermatozoa; sperm extraction and ATAC-seq of zebrafish sperm; single nuclei ATAC-seq of adult mouse testes.

### Statistical analysis and Data display

Outliers were identified using the ROUT method (Q = 1%) and removed prior to analysis. Normality was assessed using the D’Agostino–Pearson test. Depending on data distribution, either parametric or non-parametric statistical tests were applied using GraphPad Prism 10 software. For all datasets, 95% confidence intervals for mean differences were calculated using Welch’s unpaired t-test. Data are reported as differences in means for normally distributed data or differences in medians for non-normally distributed data. Percentage changes relative to control ± SE were calculated for descriptive purposes. All statistics related to omics data and heatmaps were performed using the respective packages and core R. All quantifications were carried out either blindly or by a person who had no previous knowledge of sample identities.

### Use of Artificial intelligence (AI)-assisted tools

AI-assisted tools were used during manuscript preparation and data analysis. Segmentation of zebrafish 3 dpf hearts was performed using an updated version of the deep learning pipeline described by (Ernst et al., 2023), based on convolutional neural networks with a U-Net architecture (<u>Acquifer</u>). All automated segmentation results were subsequently manually verified using visualization videos of the segmentation process.

Grammarly and ChatGPT (OpenAI) were used to improve grammar, language, and text clarity. Claude (Anthropic) and GitHub Copilot were used to assist with code development, debugging, and scripting. All AI-generated text, code, and suggestions were critically reviewed, validated, and, where necessary, modified by the authors. The authors take full responsibility for the accuracy, integrity, and final content of the manuscript, analyses, code, and segmentation results.

## Acknowledgments

We thank the animal facility, the histology and pathology unit, the advanced imaging unit, transgenesis unit, and the genomics unit of the Centro Nacional de Investigaciones Cardiovasculares CNIC. The CNIC is supported by the Instituto de Salud Carlos III (ISCIII), the Ministerio de Ciencia e Innovación (MCIN) and the Pro CNIC Foundation and is a Severo Ochoa Center of Excellence (SEV-2015-0505). We thank the Next Generation Sequencing Platform, the Core Facility for Proteomics and Mass Spectrometry and MIC-Bern at the University of Bern and Anna Gliwa, Ahmet Kürk, and Eduardo Diaz for zebrafish husbandry. We thank Suphansa Sawamiphak’s lab for sharing the *Tg(hsp70l:ifng1-2-V5, cryaa:Cerulean)* line. We thank Dr. Thierry Pedrazzini and Dr. Victor Corces as well as all the lab members of the N.M. lab for sharing knowledge and advice.

## Author contributions

Conceptualization: B.C. N.M.

Methodology: B.C., P.A., E.R., O.B.

Investigation: B.C., A.S.M., M.G., T.M., A.M.M.P., N.K., G.G., I.J.M., K.S., P.A.

Visualization: B.C., N.M., P.A.

Supervision: B.C., N.M., E.R., O.B.

Writing - original draft: B.C., N.M. P.A.

Writing - review & editing: B.C., N.M. P.A., A.S.M., A.M.M.P, T.M., N.K., G.G., I.J.M, E.R, O.B.

Funding: B.C., N.M., O.B.

