## Supplementary file for "Paternal cardiac injury elicits an inflammatory signal relay to the gonads with intergenerational cardiac effects in vertebrates"

Coppe *et al.*

**This PDF file includes:**

Figs. S1 to S14
Movies S1
Supplementary Text

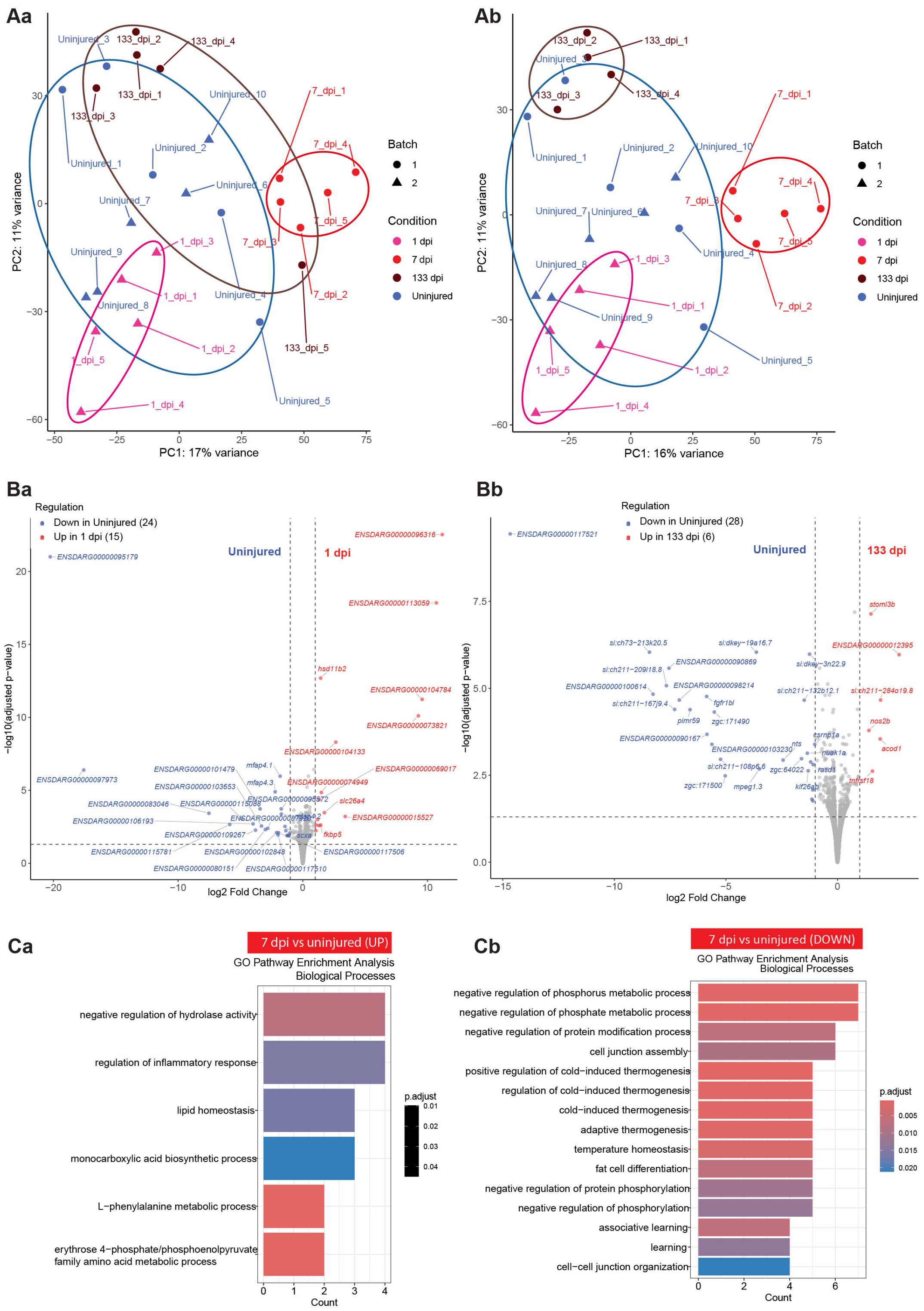


**Fig. S1. Transcriptomics analysis of zebrafish testes extracted after cardiac cryoinjury**

(A) Principal Component analysis (PCA) of RNA-seq results from uninjured testes as replicates from samples extracted at 1, 7 or 133 days postinjury (dpi). Aa shows all samples, Ab shows PCA after elimination of the sample 133_dpi_5 identified as an outlier. (B) Volcano plot showing differentially expressed genes (DEG) in testes from 1 dpi and uninjured zebrafish (Ba) and DEGs between testes samples from 133 dapi and uninjured zebrafish (Bb). Cut-off for significant DEGs was at $\log_2$ Fold change ≥ 1 and ≤ -1 and padjust ≤ 0.05. (C) Gene ontology (GO) pathway enrichment analysis of genes upregulated (Ca) or downregulated (Cb) at 7 dpi compared to uninjured testis.

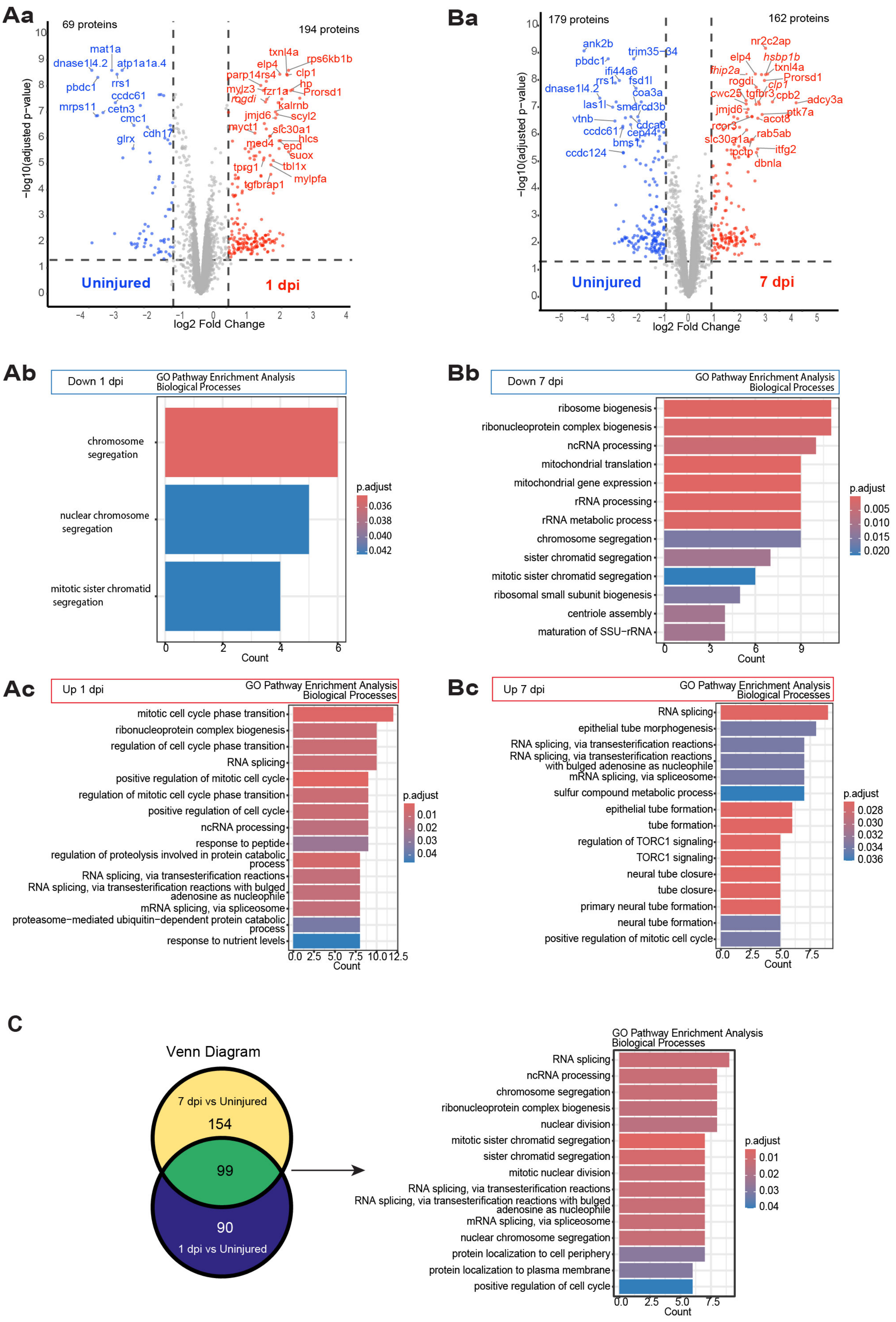
Aa
69 proteins
194 proteins
Uninjured
1 dpi
−log10(adjusted p−value)
log2 Fold Change
mat1a
dnase1l4.2
atp1a1a.4
rrs1
pbdc1
ccdc61
mrps11
cetn3
cmc1
cdh17
glrx
txnl4a
rps6kb1b
elp4
clp1
parp14rs4
hp
mylz3
fzr1a
Prorsd1
rogdi
kalrnb
jmjd6
scyl2
myct1
slc30a1
hlcs
med4
epd
suox
tprg1
tbl1x
mylpfa
tgfbrap1
Ba
179 proteins
162 proteins
Uninjured
7 dpi
−log10(adjusted p−value)
log2 Fold Change
ank2b
trim35−34
pbdc1
ifi44a6
rrs1
fsd1l
dnase1l4.2
coa3a
las1l
smarcd3b
vtnb
cdca8
ccdc61
cep44
bms1
ccdc124
nr2c2ap
elp4
hsbp1b
txnl4a
thip2a
Prorsd1
rogdi
clp1
cwc25
tgfbr3
cpb2
adcy3a
jmjd6
ptk7a
acot8
rcor3
rab5ab
slc30a1a
pctp
itfg2
dbnla
Ab
Down 1 dpi
GO Pathway Enrichment Analysis
Biological Processes
chromosome segregation
nuclear chromosome segregation
mitotic sister chromatid segregation
Count
p.adjust
Bb
Down 7 dpi
GO Pathway Enrichment Analysis
Biological Processes
ribosome biogenesis
ribonucleoprotein complex biogenesis
ncRNA processing
mitochondrial translation
mitochondrial gene expression
rRNA processing
rRNA metabolic process
chromosome segregation
sister chromatid segregation
mitotic sister chromatid segregation
ribosomal small subunit biogenesis
centriole assembly
maturation of SSU−rRNA
Count
p.adjust
Ac
Up 1 dpi
GO Pathway Enrichment Analysis
Biological Processes
mitotic cell cycle phase transition
ribonucleoprotein complex biogenesis
regulation of cell cycle phase transition
RNA splicing
positive regulation of mitotic cell cycle
regulation of mitotic cell cycle phase transition
positive regulation of cell cycle
ncRNA processing
response to peptide
regulation of proteolysis involved in protein catabolic process
RNA splicing, via transesterification reactions
RNA splicing, via transesterification reactions with bulged adenosine as nucleophile
mRNA splicing, via spliceosome
proteasome−mediated ubiquitin−dependent protein catabolic process
response to nutrient levels
Count
p.adjust
Bc
Up 7 dpi
GO Pathway Enrichment Analysis
Biological Processes
RNA splicing
epithelial tube morphogenesis
RNA splicing, via transesterification reactions
RNA splicing, via transesterification reactions with bulged adenosine as nucleophile
mRNA splicing, via spliceosome
sulfur compound metabolic process
epithelial tube formation
tube formation
regulation of TORC1 signaling
TORC1 signaling
neural tube closure
tube closure
primary neural tube formation
neural tube formation
positive regulation of mitotic cell cycle
Count
p.adjust
C
Venn Diagram
7 dpi vs Uninjured
154
99
90
1 dpi vs Uninjured
GO Pathway Enrichment Analysis
Biological Processes
RNA splicing
ncRNA processing
chromosome segregation
ribonucleoprotein complex biogenesis
nuclear division
mitotic sister chromatid segregation
sister chromatid segregation
mitotic nuclear division
RNA splicing, via transesterification reactions
RNA splicing, via transesterification reactions with bulged adenosine as nucleophile
mRNA splicing, via spliceosome
nuclear chromosome segregation
protein localization to cell periphery
protein localization to plasma membrane
positive regulation of cell cycle
Count
p.adjust

**Fig. S2. Proteomics analysis of testes extracted from zebrafish after cardiac cryoinjury.**
**(A)** Comparison of testes at 1 day postinjury (dpi) vs uninjured testes. Zebrafish proteins were converted into mouse gene orthologs. Shown is the volcano plot (**Aa**), and Gene Ontology (GO) pathway analysis of differentially downregulated (**Ab**) and upregulated (**Ac**) proteins. **(B)** Comparison of testes at 7dpi vs uninjured testes. Zebrafish proteins were converted into mouse gene orthologs. Shown is the volcano plot (**Ba**), and Gene Ontology (GO) pathway analysis of differentially downregulated (**Bb**) and upregulated (**Bc**) proteins. Criteria for differential expression were $log_2$ fold change≥1 and ≤-1 and p-adjusted value ≤ 0-05 for data shown in A and B. **(C)** Venn Diagram revealing common and time point specific protein changes (left) and GO pathway analysis of common proteins between time points (right).

A

B

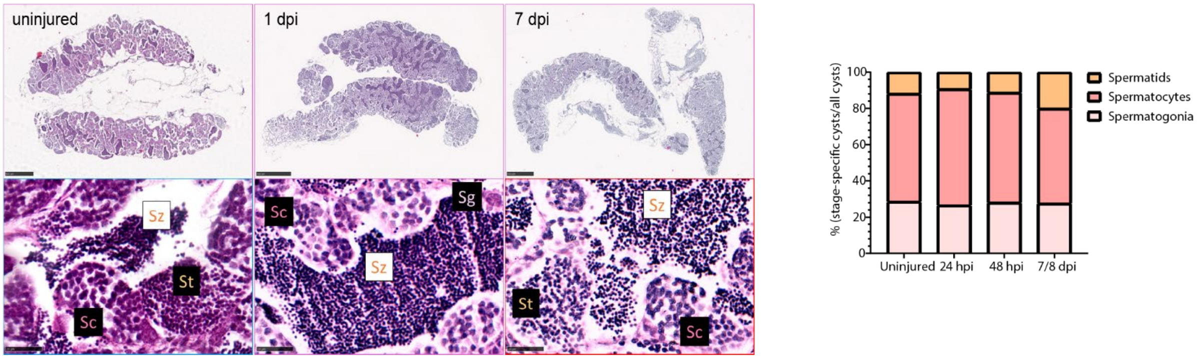


**Fig. S3. Histology of testes and analysis of sperm maturation after cardiac cryoinjury.**
(**A)** Hematoxylin&Eosin histological stainings of uninjured testis and testis collected at 1 and 7 dpi. Scale bar: 500 µm. Lower rows represent magnified views of cysts. Scale bar: 25µm. (**B)** Classification of the percentage of each cell type relative to the total number of cysts per seminiferous tubule at the indicated condition. For each animal, 2-3 seminiferous tubules were analyzed, with 5-6 animals included per experimental group. Statistical analysis: Tukey's multiple comparisons test. Dpi: days post-injury; Sc: spermatocytes; Sg: spermatogonia; St: spermatids; Sz: spermatozoa.

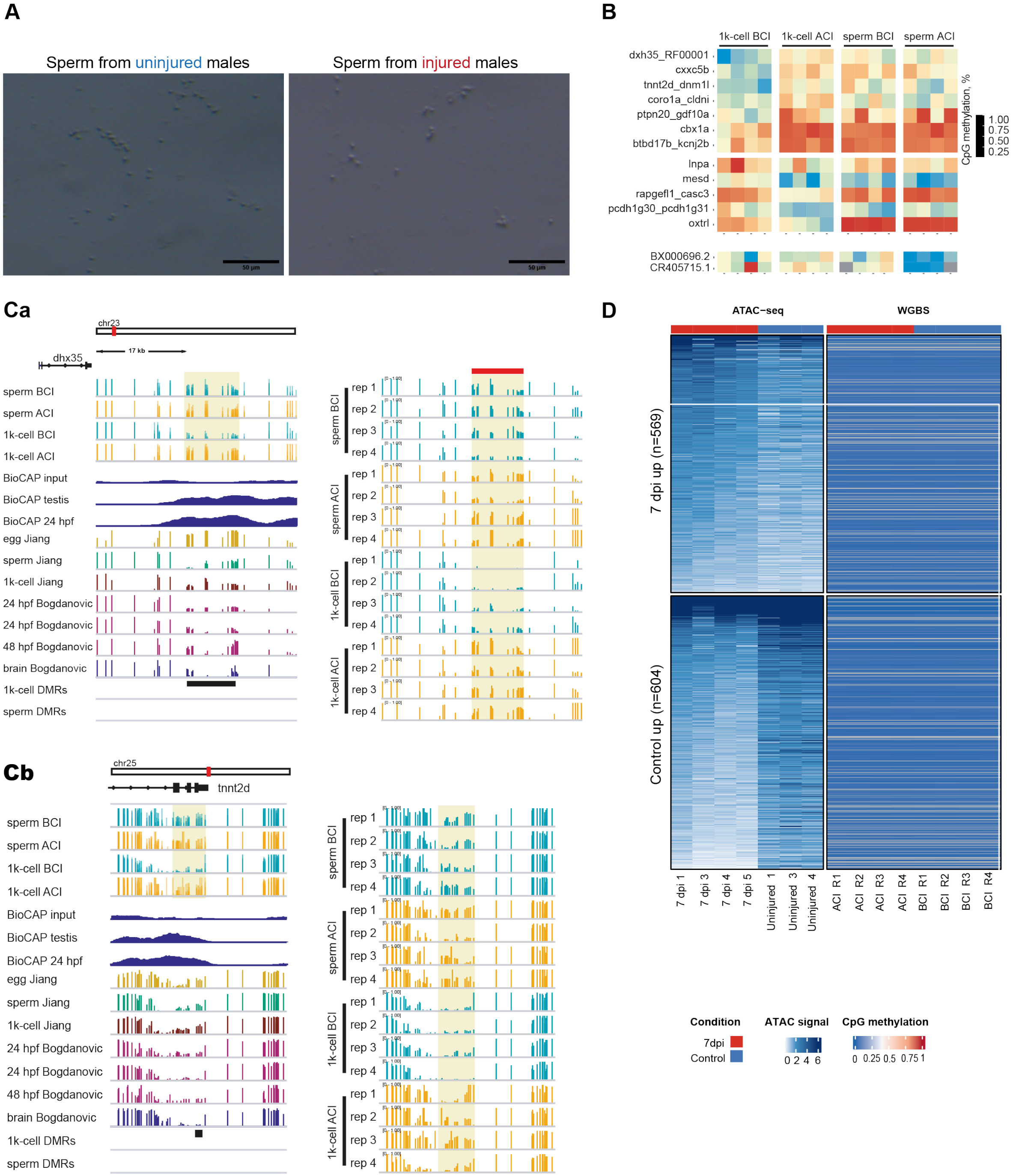


**Fig. S4. Assessment of DNA methylation changes in sperm and F1 zebrafish embryos as well as chromatin accessibility changes in sperm after cardiac cryoinjury.**

**(A)** Images of freshly extracted sperm from injured (after cryo-injury: ACI) and uninjured (before cryo-injury: BCI) zebrafish. (**B**) Heat map of differentially methylated regions (DMRs) found in samples from F1 embryos from crosses of uninjured males (1k-cell stage BCI) or from crosses of males at 7dpi (1k-cell stage ACI), as well as sperm samples collected two weeks before or seven days after cardiac cryoinjury. (**C)** Here are shown the two DMR **(***dhx35* (upper panel) and *tnnt2d* (lower panel)) found in F1 from injured (7 dpi) and uninjured father at the 1k-cell stage in comparison with methylome databases of other repositories at different time points. (**D)** Heatmap of ATAC-seq and Whole Bisulfite Genome sequencing (WGBS) of the different replicates of 1k-cell stage embryos and sperm as explained in B. Note that there are differences observed in chromatin accessibility between uninjured (blue) and injured (red) conditions, but no differences were found for WGBS analysis.

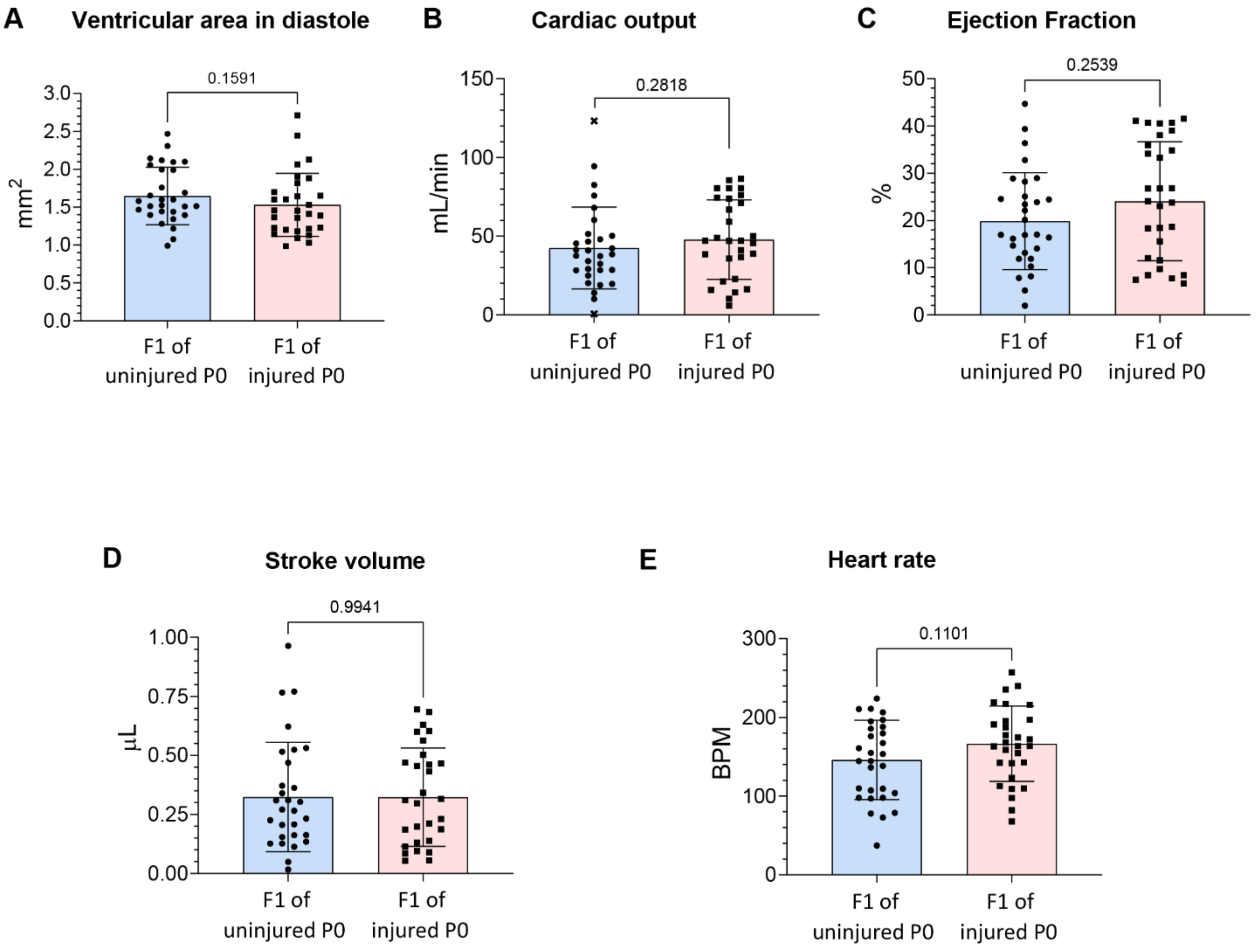


**Fig. S5. Echocardiographic measurements of adult F1 from injured or uninjured zebrafish males.**

**(A-E)** Shown are different cardiac function parameters as indicated above the graphs. Shown are measurements of individual F1 animals as well as mean and standard deviation. Statistical test employed: (A-D) Mann Whitey test, (E), unpaired T test.

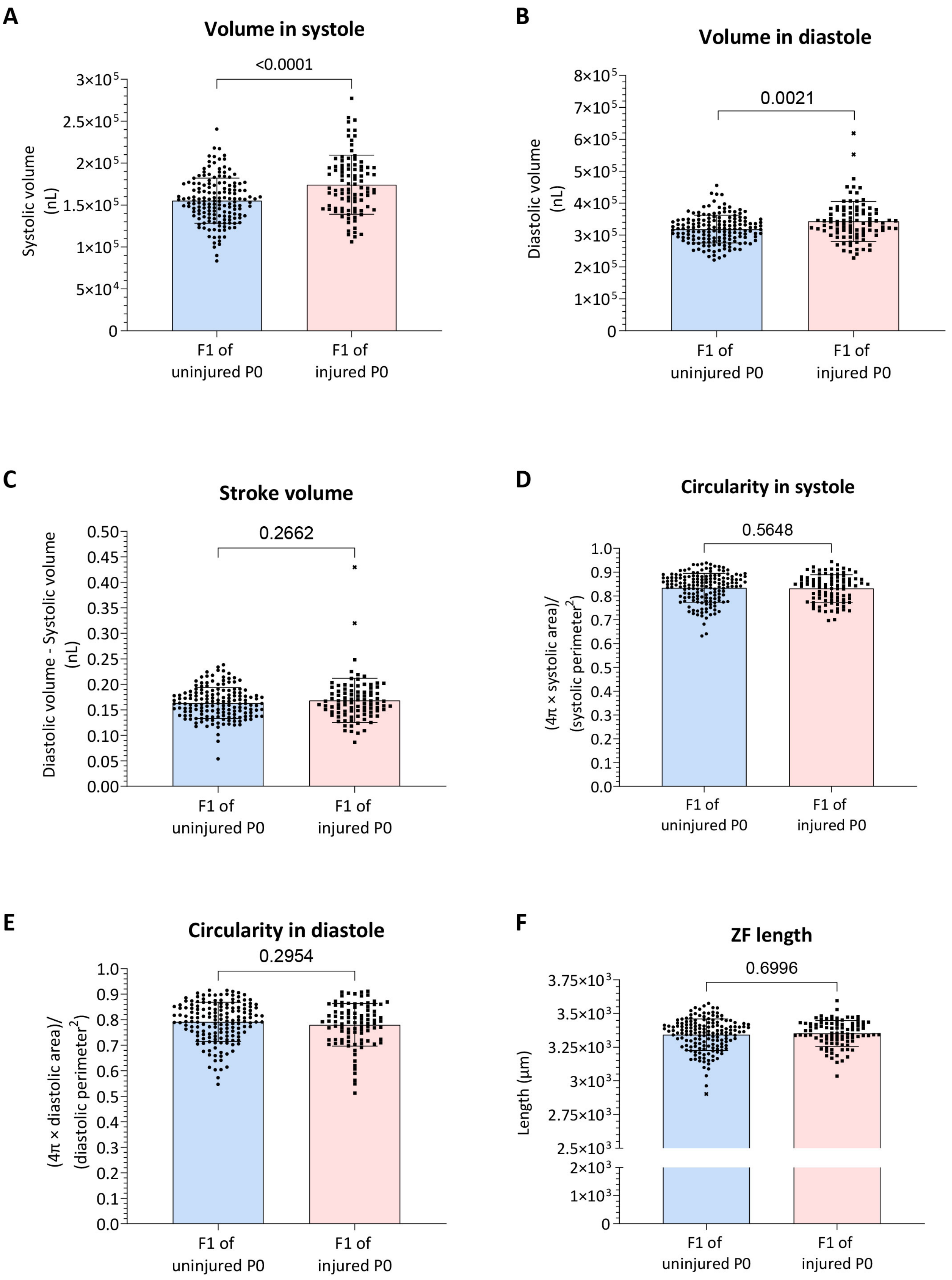
A
Volume in systole
<0.0001
Systolic volume (nL)
3×10^5
2.5×10^5
2×10^5
1.5×10^5
1×10^5
5×10^4
0
F1 of uninjured P0
F1 of injured P0
B
Volume in diastole
0.0021
Diastolic volume (nL)
8×10^5
7×10^5
6×10^5
5×10^5
4×10^5
3×10^5
2×10^5
1×10^5
0
F1 of uninjured P0
F1 of injured P0
C
Stroke volume
0.2662
Diastolic volume - Systolic volume (nL)
0.50
0.45
0.40
0.35
0.30
0.25
0.20
0.15
0.10
0.05
0.00
F1 of uninjured P0
F1 of injured P0
D
Circularity in systole
0.5648
(4π × systolic area)/ (systolic perimeter^2)
1.0
0.9
0.8
0.7
0.6
0.5
0.4
0.3
0.2
0.1
0.0
F1 of uninjured P0
F1 of injured P0
E
Circularity in diastole
0.2954
(4π × diastolic area)/ (diastolic perimeter^2)
1.0
0.9
0.8
0.7
0.6
0.5
0.4
0.3
0.2
0.1
0.0
F1 of uninjured P0
F1 of injured P0
F
ZF length
0.6996
Length (μm)
3.75×10^3
3.5×10^3
3.25×10^3
3×10^3
2.75×10^3
2.5×10^3
2×10^3
1×10^3
0
F1 of uninjured P0
F1 of injured P0

**Fig. S6. Cardiac function assessment of larval F1 hearts from crosses with injured or uninjured zebrafish males.**

**(A-F)** Shown are different cardiac function parameters as indicated above the graphs. Shown are measurements of individual F1 animals at 3 dpf as well as mean and standard deviation. Statistical test employed: Unpaired t test (A-C); Mann Whitey test (D-F).

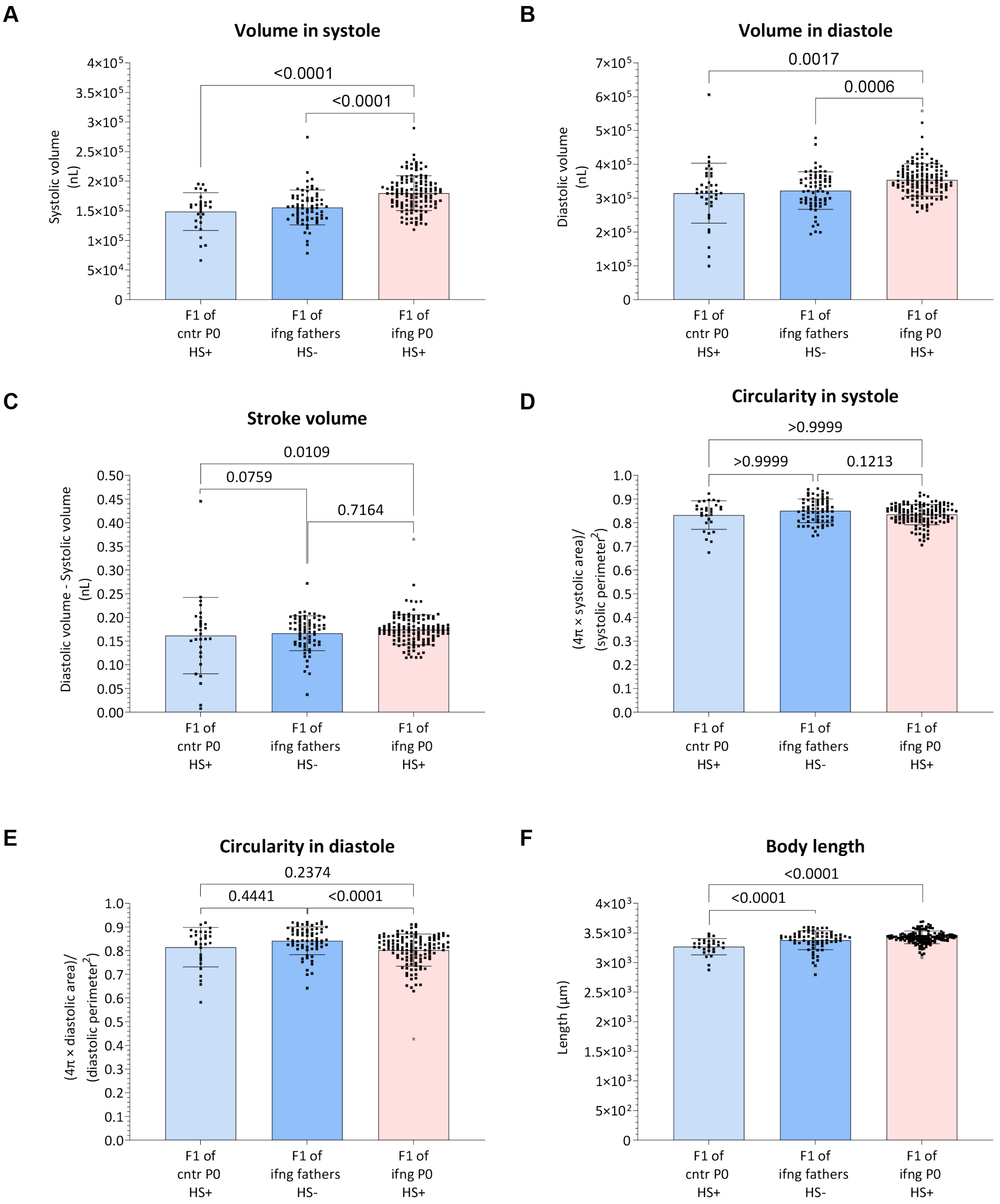


Fig. S7. Cardiac function assessment of larval F1 hearts from crosses with zebrafish males that underwent interferon gamma systemic overexpression.

Adult *hsp70:ifng1* males underwent one heat shock and were crossed with wildtype (WT) females after one week. The larvae were collected and cardiac function assessed at 3 days postfertilization (dpf). Control larvae were obtained from crosses of females with non-heat-shocked *hsp70:ifng1* males (F1 of ifng father HS-) and additionally from crosses of heat-shocked WT males (F1 of cntr P0 HS+). Shown are different cardiac function parameters as indicated above the graphs. Shown are measurements of individual F1 animals at 3 dpf as well as mean and standard deviation. Statistical test employed: Ordinary one-way ANOVA, Turkey's multiple comparisons test (A, C); Kruskal-Wallis, Dunn's multiple comparisons test (B,D,E,F)

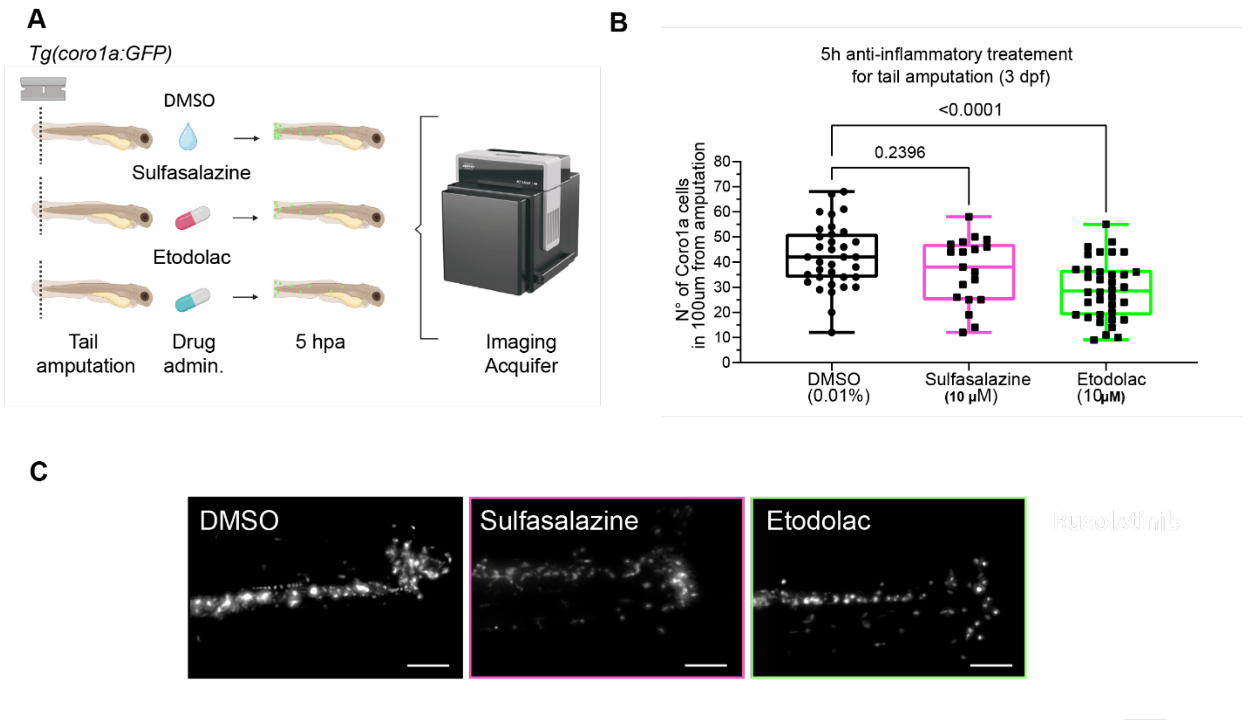


**Fig. S8. Effect of anti-inflammatory compounds on leukocyte migration**

**(A)** Tails of *Tg(coro1a:GFP)* larvae were amputated and DMSO, Sulfasalazine or Etodolac administered to the larvae in water. After five hours postamputation (hpa), *coro1a*:GFP-positive leukocytes at the amputation plane were counted. Image created in BioRender. Coppe, B. (2026) https://BioRender.com/6m9279f **(B)** Graph showing the number of *coro1a*: GFP-positive cells present at the 100 µm amputation border zone in the three conditions. Each dot represents number of cells in one larva. Shown are box blot and minimum to maximum. Statistics: Ordinary One way Anova, Turkey's multiple comparisons test. **(C)** Representative images of the tail amputation region (posterior end to the right) showing *coro1a*:GFP-positive cells accumulation at the amputation plane. Note: lower amount of cells upon Etodolac treatment compared to DMSO and Sulfosalazine treated larvae. Scale bar: 100 µm.

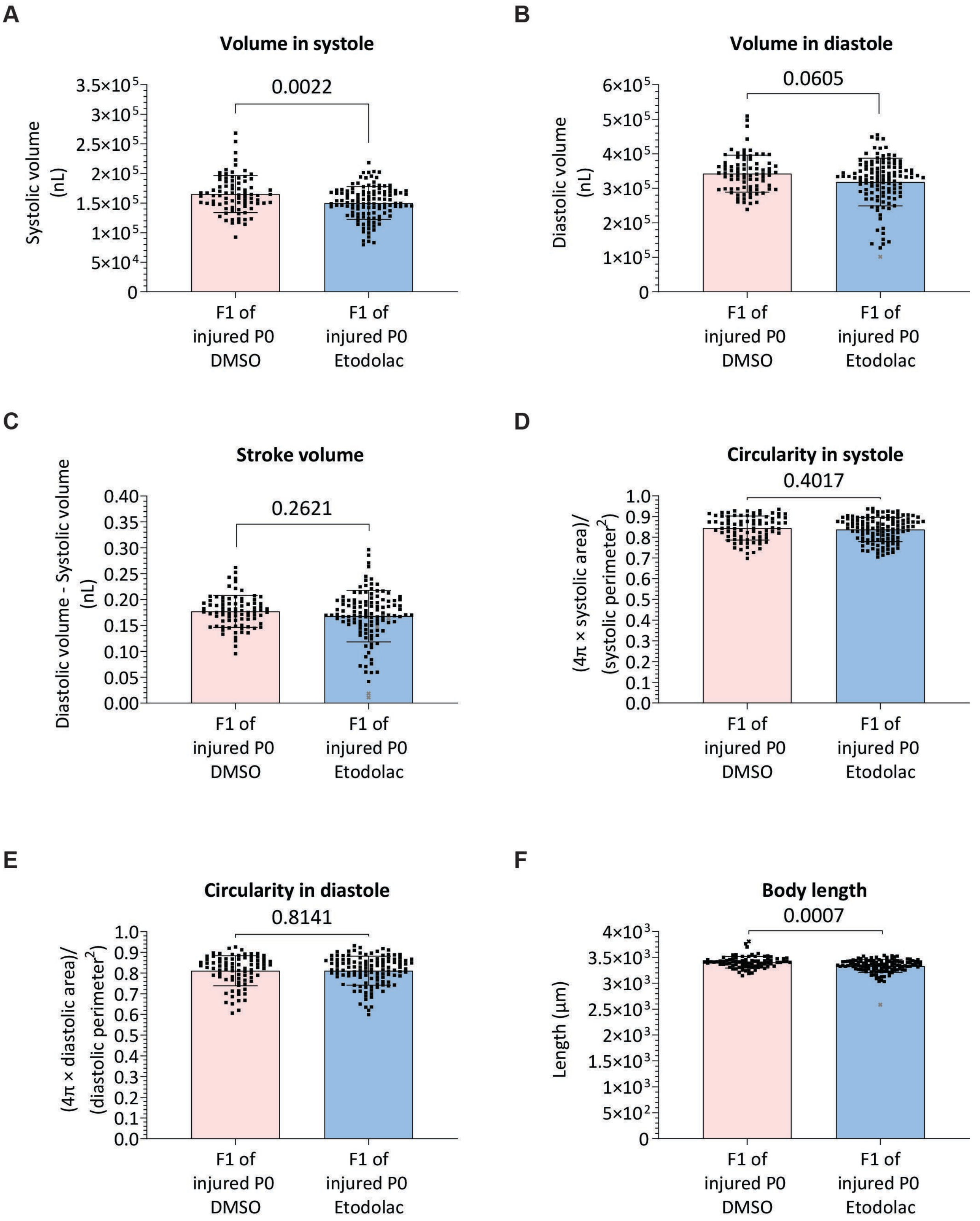


**Fig S9. Cardiac function assessment of larval F1 hearts from crosses with injured zebrafish males that underwent anti-inflammatory treatment.** Adult wild-type (AB) males underwent cardiac cryoinjury followed by daily DMSO or Etodolac injection for one week. Then, they were crossed with *Tg(myl7:eGFP)*

females. The larvae were collected and cardiac function assessed at 3 days postfertilization (dpf). Shown are different cardiac function parameters as indicated above the graphs. Shown are measurements of individual F1 animals as well as mean and standard deviation. Statistical test employed: Mann-Whitney test (A, B,D,E,F); Unpaired t test (C).

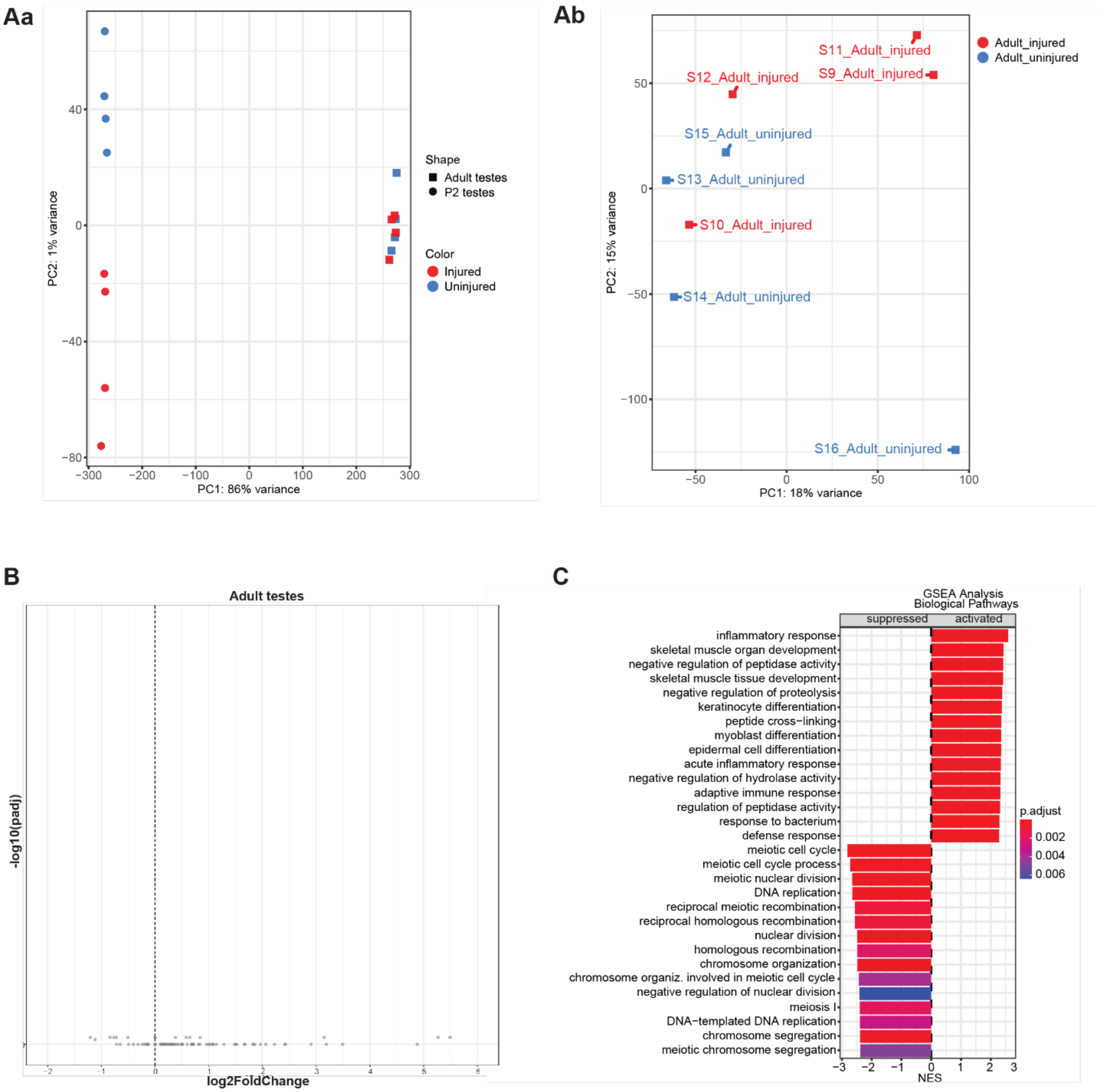


**Fig. S10. Transcriptomic analysis of testis (neonatal and adult) from injured and uninjured male mice.**

(A) PCA plots of samples from P2 and adult testis collected from injured (CI at P1) and uninjured mouse males. (Aa) combined PCA of both states, P2 and adult. (Ab) PCA plot of only adult samples. (B) Volcano plot of adult testis RNA-seq. Note that there are no gene expression changes between adult gonads from injured (CI at P1) and uninjured males. (C) Gene set enrichment analysis of RNA-seq of P2 testis of injured (CI at P1) and uninjured males.

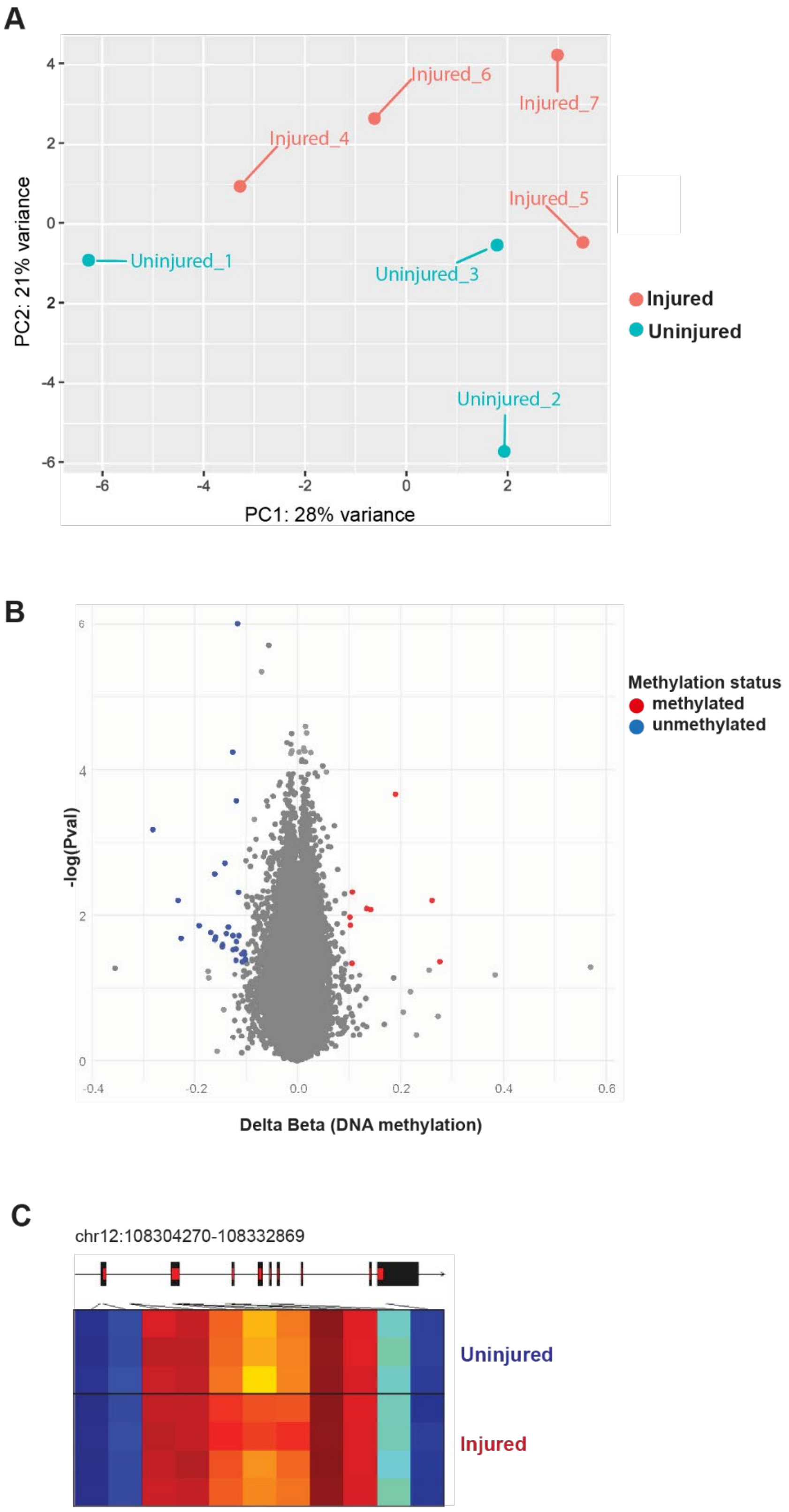
A
Injured_6
Injured_7
Injured_4
Injured_5
Uninjured_1
Uninjured_3
Uninjured_2
PC2: 21% variance
PC1: 28% variance
Injured
Uninjured
B
Methylation status
methylated
unmethylated
-log(Pval)
Delta Beta (DNA methylation)
C
chr12:108304270-108332869
Uninjured
Injured

**Fig. S11. DNA Methylation analysis in sperm of adult injured (CI performed at P1) and uninjured mice.** (**A**) Principal Component Analysis (PCA) of DNA methylation *loci* in testis samples from male mice that were cryoinjured at P1 and testis samples from uninjured control mice. (**B**) Volcano plot of differentially methylated loci (i.e. probes) in testis from injured and uninjured mice. Methylation levels are expressed in colors, where blue corresponds to unmethylated and red to methylated loci. Differentially methylated probes were filtered by p-value (≤0.05) and effect size (≥10%). (**C**) Example of a differentially methylated genomic locus in sperm from injured or uninjured male. Each column represents a probe/locus, each row one sample. Methylation levels are shown in colors, based on a blue-red scale where blue corresponds to unmethylated and red to fully methylated DNA. No differentially methylated regions (DMRs) were identified in sperm from injured *vs* uninjured mice.

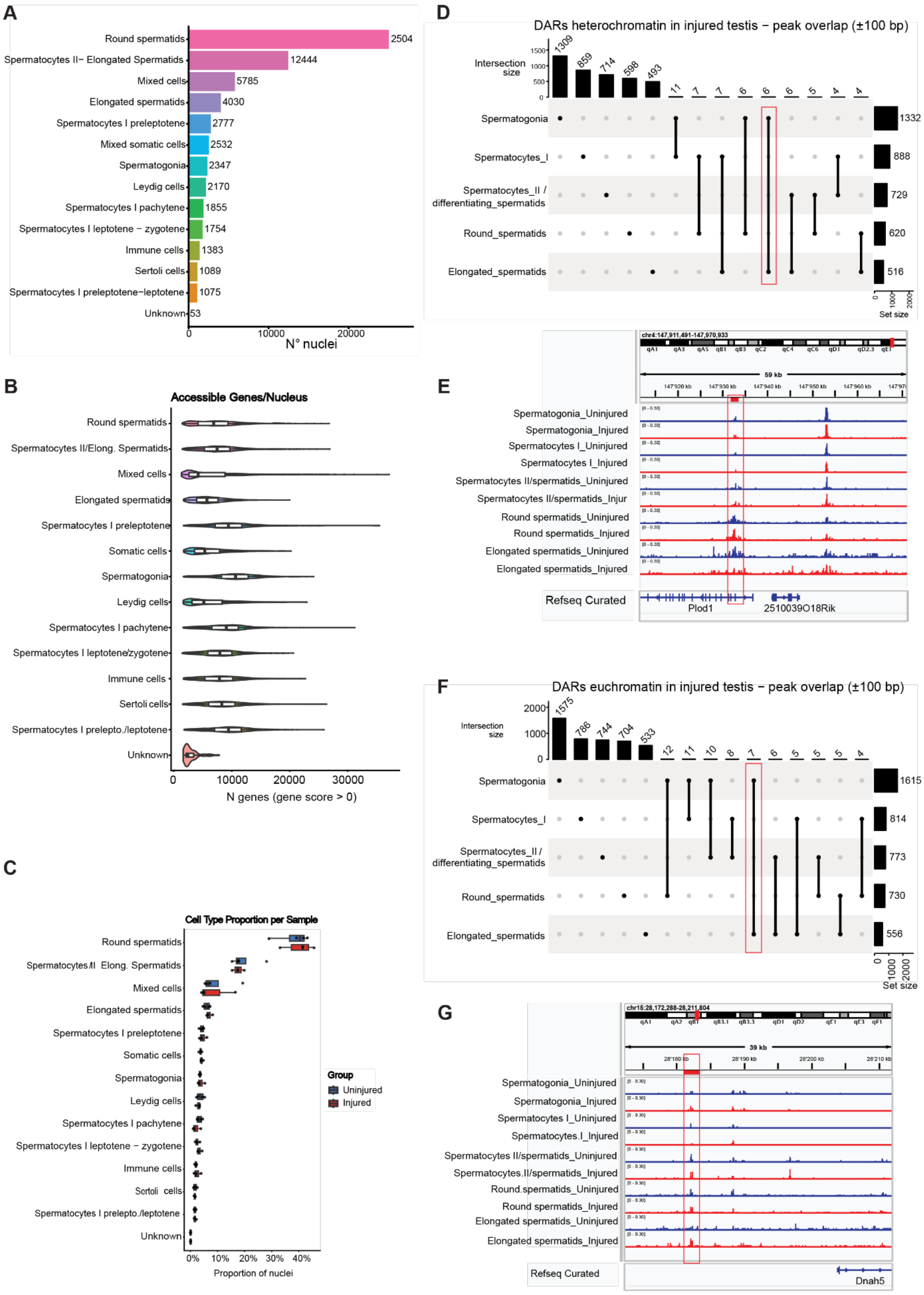
A
Round spermatids 2504
Spermatocytes II– Elongated Spermatids 12444
Mixed cells 5785
Elongated spermatids 4030
Spermatocytes I preleptotene 2777
Mixed somatic cells 2532
Spermatogonia 2347
Leydig cells 2170
Spermatocytes I pachytene 1855
Spermatocytes I leptotene – zygotene 1754
Immune cells 1383
Sertoli cells 1089
Spermatocytes I preleptotene–leptotene 1075
Unknown 53
N° nuclei
B
Accessible Genes/Nucleus
N genes (gene score > 0)
C
Cell Type Proportion per Sample
Group
Uninjured
Injured
Proportion of nuclei
D
DARs heterochromatin in injured testis – peak overlap (±100 bp)
Intersection size
Spermatogonia 1332
Spermatocytes_I 888
Spermatocytes_II / differentiating_spermatids 729
Round_spermatids 620
Elongated_spermatids 516
Set size
E
Spermatogonia_Uninjured
Spermatogonia_Injured
Spermatocytes I_Uninjured
Spermatocytes I_Injured
Spermatocytes II/spermatids_Uninjured
Spermatocytes II/spermatids_Injur
Round spermatids_Uninjured
Round spermatids_Injured
Elongated spermatids_Uninjured
Elongated spermatids_Injured
Refseq Curated
Plod1
2510039O18Rik
F
DARs euchromatin in injured testis – peak overlap (±100 bp)
Intersection size
Spermatogonia 1615
Spermatocytes_I 814
Spermatocytes_II / differentiating_spermatids 773
Round_spermatids 730
Elongated_spermatids 556
Set size
G
Spermatogonia_Uninjured
Spermatogonia_Injured
Spermatocytes I_Uninjured
Spermatocytes.I_Injured
Spermatocytes II/spermatids_Uninjured
Spermatocytes.II/spermatids_Injured
Round.spermatids_Uninjured
Round spermatids_Injured
Elongated spermatids_Uninjured
Elongated spermatids_Injured
Refseq Curated
Dnah5

**Fig. S12. Single-nuclei chromatin accessibility assay of testes collected from adult male mice with and without a cardiac cryoinjury (performed at P1).**

(**A**) Overall number of nuclei obtained from different gonadal cell types. (**B**) Number of accessible genes per nucleus detected in different gonadal cell types. (**C**) Proportion of cell types from which cell nuclei were obtained in the two experimental groups. (**D**) UpSet plot displaying common regions within gonadal cell types that are in a heterochromatic state (not accessible, closed regions) in samples from injured animals compared to uninjured animals. Highlighted are common DARs in spermatogonia and elongated spermatids. (**E**) Example of a region containing a DAR in heterochromatin state in both spermatogonia and elongated spermatids of injured samples. (**F**) UpSet plot displaying regions within gonadal cell types that are in a euchromatic state (accessible, open regions) in samples from injured animals compared to uninjured animals. Highlighted are common DARs in spermatogonia and elongated spermatids. (**G**) Example region of a peak that is found in euchromatin state in both spermatogonia and elongated spermatids of injured samples.

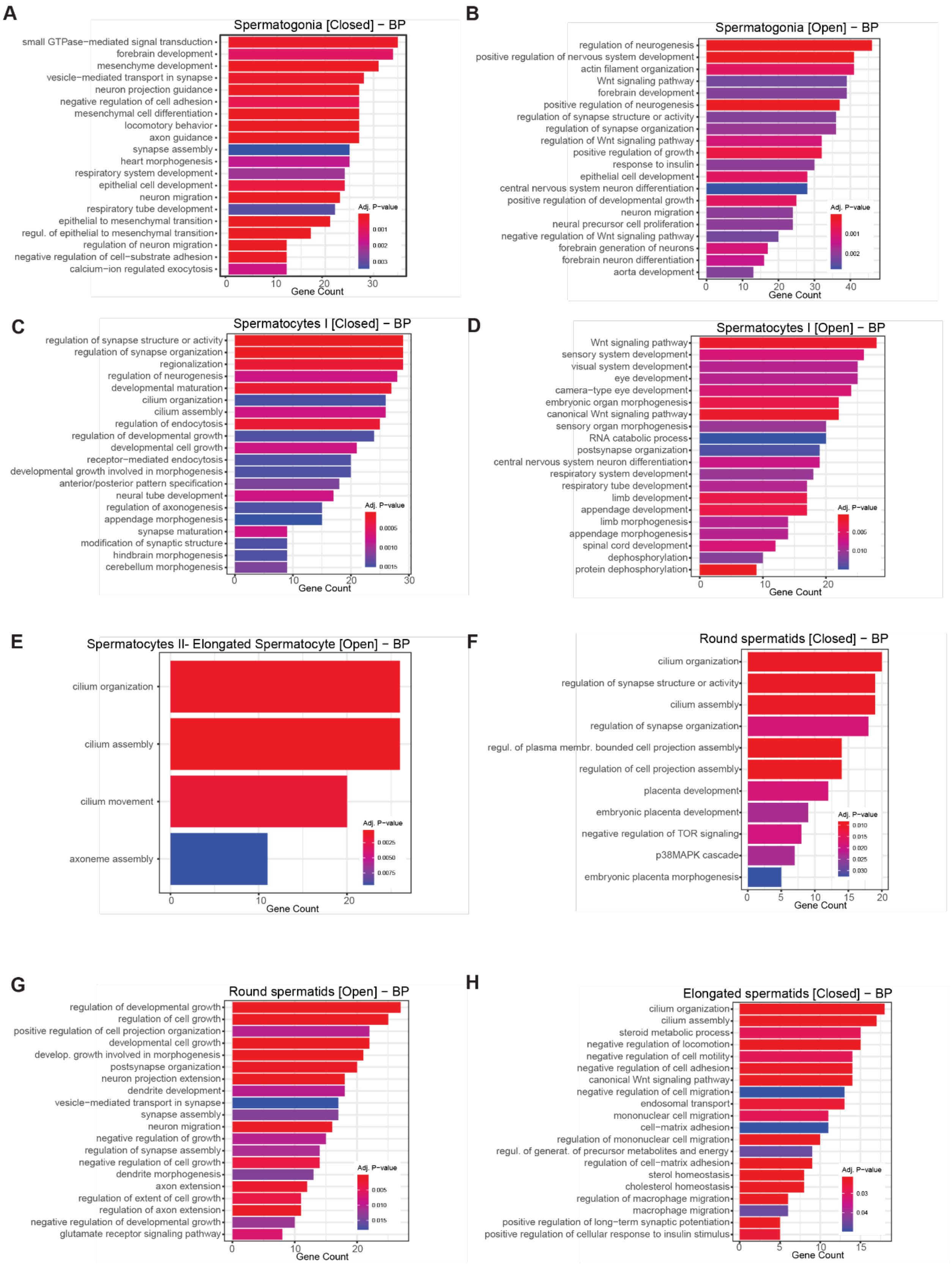
A
Spermatogonia [Closed] − BP
small GTPase−mediated signal transduction
forebrain development
mesenchyme development
vesicle−mediated transport in synapse
neuron projection guidance
negative regulation of cell adhesion
mesenchymal cell differentiation
locomotory behavior
axon guidance
synapse assembly
heart morphogenesis
respiratory system development
epithelial cell development
neuron migration
respiratory tube development
epithelial to mesenchymal transition
regul. of epithelial to mesenchymal transition
regulation of neuron migration
negative regulation of cell−substrate adhesion
calcium−ion regulated exocytosis
Adj. P−value
0.001
0.002
0.003
0
10
20
30
Gene Count
B
Spermatogonia [Open] − BP
regulation of neurogenesis
positive regulation of nervous system development
actin filament organization
Wnt signaling pathway
forebrain development
positive regulation of neurogenesis
regulation of synapse structure or activity
regulation of synapse organization
regulation of Wnt signaling pathway
positive regulation of growth
response to insulin
epithelial cell development
central nervous system neuron differentiation
positive regulation of developmental growth
neuron migration
neural precursor cell proliferation
negative regulation of Wnt signaling pathway
forebrain generation of neurons
forebrain neuron differentiation
aorta development
Adj. P−value
0.001
0.002
0
10
20
30
40
Gene Count
C
Spermatocytes I [Closed] − BP
regulation of synapse structure or activity
regulation of synapse organization
regionalization
regulation of neurogenesis
developmental maturation
cilium organization
cilium assembly
regulation of endocytosis
regulation of developmental growth
developmental cell growth
receptor−mediated endocytosis
developmental growth involved in morphogenesis
anterior/posterior pattern specification
neural tube development
regulation of axonogenesis
appendage morphogenesis
synapse maturation
modification of synaptic structure
hindbrain morphogenesis
cerebellum morphogenesis
Adj. P−value
0.0005
0.0010
0.0015
0
10
20
30
Gene Count
D
Spermatocytes I [Open] − BP
Wnt signaling pathway
sensory system development
visual system development
eye development
camera−type eye development
embryonic organ morphogenesis
canonical Wnt signaling pathway
sensory organ morphogenesis
RNA catabolic process
postsynapse organization
central nervous system neuron differentiation
respiratory system development
respiratory tube development
limb development
appendage development
limb morphogenesis
appendage morphogenesis
spinal cord development
dephosphorylation
protein dephosphorylation
Adj. P−value
0.005
0.010
0
10
20
Gene Count
E
Spermatocytes II- Elongated Spermatocyte [Open] − BP
cilium organization
cilium assembly
cilium movement
axoneme assembly
Adj. P−value
0.0025
0.0050
0.0075
0
10
20
Gene Count
F
Round spermatids [Closed] − BP
cilium organization
regulation of synapse structure or activity
cilium assembly
regulation of synapse organization
regul. of plasma membr. bounded cell projection assembly
regulation of cell projection assembly
placenta development
embryonic placenta development
negative regulation of TOR signaling
p38MAPK cascade
embryonic placenta morphogenesis
Adj. P−value
0.010
0.015
0.020
0.025
0.030
0
5
10
15
20
Gene Count
G
Round spermatids [Open] − BP
regulation of developmental growth
regulation of cell growth
positive regulation of cell projection organization
developmental cell growth
develop. growth involved in morphogenesis
postsynapse organization
neuron projection extension
dendrite development
vesicle−mediated transport in synapse
synapse assembly
neuron migration
negative regulation of growth
regulation of synapse assembly
negative regulation of cell growth
dendrite morphogenesis
axon extension
regulation of extent of cell growth
regulation of axon extension
negative regulation of developmental growth
glutamate receptor signaling pathway
Adj. P−value
0.005
0.010
0.015
0
10
20
Gene Count
H
Elongated spermatids [Closed] − BP
cilium organization
cilium assembly
steroid metabolic process
negative regulation of locomotion
negative regulation of cell motility
negative regulation of cell adhesion
canonical Wnt signaling pathway
negative regulation of cell migration
endosomal transport
mononuclear cell migration
cell−matrix adhesion
regulation of mononuclear cell migration
regul. of generat. of precursor metabolites and energy
regulation of cell−matrix adhesion
sterol homeostasis
cholesterol homeostasis
regulation of macrophage migration
macrophage migration
positive regulation of long−term synaptic potentiation
positive regulation of cellular response to insulin stimulus
Adj. P−value
0.03
0.04
0
5
10
15
Gene Count

**Fig. S13. Pathway analysis of genes associated to differential accessible regions in the germ line of injured versus uninjured male mice. (A-H)** Shown are gene ontology biological pathway (BP) analysis of genes neighboring open or closed chromatin regions. Panels A-H show data for different germ cell stages.

A

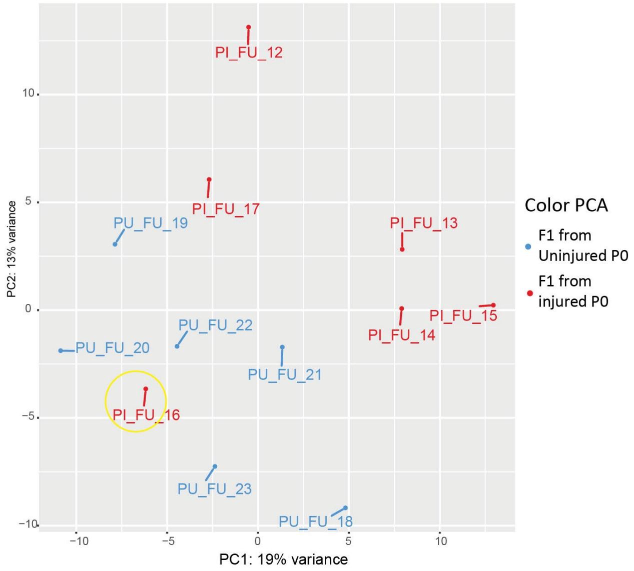


B

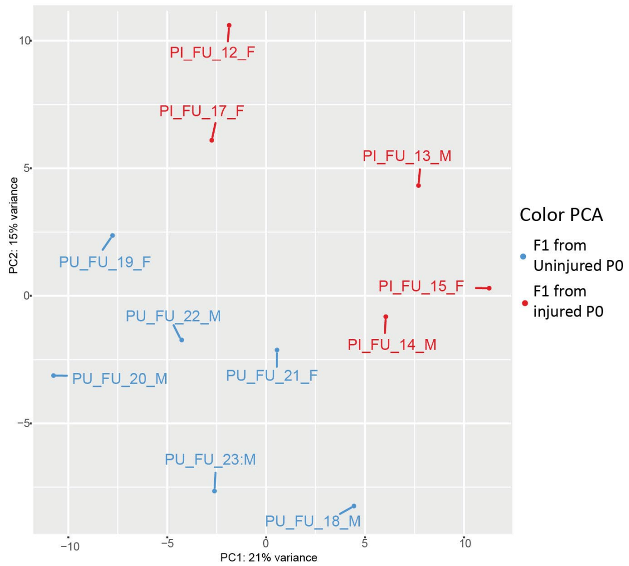


**Fig. S14. Principal component analysis (PCA) of transcriptomic analysis of adult F1 mouse hearts of injured and uninjured parental generation.** (**A**) PCA plot including all biological replicates. The yellow circle marks an outlier sample. (**B**) PCA plot of the samples excluding the outlier highlighted in (A).

### Supplementary Materials and Methods

**RNA extraction and bulk RNA-seq of zebrafish and mice P0 testes and F1 hearts**

Testes from four sibling male zebrafish were pooled for each biological replicate. Five biological replicates were collected per condition (1, 7, and 133 dpi, and uninjured controls). Samples were processed in two experimental batches: experiment 1 included uninjured, 7 dpi, and 133 dpi samples, whereas experiment 2 included uninjured and 1 dpi samples. Data from both experiments were combined following batch correction. Principal component analysis (PCA) was used for quality assessment, and sample "133_dpi_5" was identified as an outlier and excluded from downstream analyses (Figure 1B; Supplementary Figure 1A).

Testes from mice were collected either one day after injury (P2) or from adult mice approximately eight months after birth. Age-matched uninjured animals served as controls. Each biological replicate consisted of a single testis.

Hearts (*i.e.* ventricles) were collected from adult zebrafish offspring of uninjured or injured males (crossed at 7 dpi). The bulbus arteriosus and atrium were removed manually, and a small incision was made in each ventricle to remove erythrocytes. Four biological replicates per condition were generated, each consisting of pooled ventricles from four sibling fish.

In mice, hearts were collected at 3 wpp from adult offspring of uninjured or injured males (CI performed at P1). Samples included four control and three experimental males, and two control and three experimental females. PCA was used for quality assessment, and sample "Injured P0_16" was identified as an outlier and excluded from downstream analyses (Figures S14 A,B).

To prepare the samples, tissues were first washed in PBS and snap frozen. On the day of RNA extraction, samples were homogenized in TRI Reagent (Sigma-Aldrich) using 21G and 30G syringes (BD Microlance). Following phase separation with 1-bromo-3-chloropropane (Sigma-Aldrich), the aqueous phase was mixed with an equal volume of ethanol, and RNA was purified using the RNA Clean & Concentrator kit (Zymo Research) with on-column DNase I treatment. RNA quantity and quality were assessed using a NanoDrop and a Fragment Analyzer before library preparation and sequencing.

Only for zebrafish F1 adult hearts RNA was extracted using the PicoPure RNA Isolation Kit (Arcturus, Thermo Fisher Scientific; Cat. Nos. KIT0202/KIT0204) following the manufacturer's instructions. Briefly, heart tissue was lysed in Extraction Buffer (XB) and incubated at 42°C for 45 min, followed by homogenization through a 30-gauge needle and centrifugation to remove debris. The clarified lysate was mixed with 70% ethanol and applied to a preconditioned purification column. After washing, on-column DNase I digestion was performed for 15 min at room temperature, followed by additional wash steps. RNA

was eluted in 11 µL Elution Buffer, quantified using a NanoDrop spectrophotometer (Thermo Fisher Scientific), and stored at −80°C until use.
Library preparation and sequencing of zebrafish testes were performed by the Next Generation Sequencing (NGS) Platform of the University of Bern (Switzerland). The quantity and quality of purified total RNA were assessed using a Qubit 4.0 fluorometer with the Qubit RNA BR Assay Kit (Thermo Fisher Scientific) and an Advanced Analytical Fragment Analyzer System with a Fragment Analyzer RNA Kit (Agilent), respectively. Libraries were prepared using the TruSeq Stranded mRNA Library Prep Kit (Illumina, 20020595) with 1 µg total RNA input and TruSeq RNA UD Indexes (Illumina, 20022371) according to the manufacturer's instructions. Resulting libraries were evaluated using Qubit dsDNA HS Assay and Fragment Analyzer HS NGS Fragment Kit. Equimolar pooled libraries were sequenced paired-end on an Illumina NovaSeq 6000 instrument using either a 100-cycle SP Reagent Kit v1.5 (2 × 51 bp reads) or a 300-cycle SP Reagent Kit (2 × 151 bp reads). Sequencing quality was assessed using Illumina Sequencing Analysis Viewer, and base call files were demultiplexed and converted into FASTQ files using Illumina bcl2fastq conversion software v2.20.

Library preparation and sequencing of zebrafish hearts and mouse samples (both hearts and testes) were performed by the Genomics Unit at CNIC (Madrid, Spain). 75 ng of RNA from zebrafish hearts was used to generate barcoded RNA-seq libraries using the Ovation Single Cell RNA-Seq System (NuGEN) with two rounds of library amplification. Zebrafish heart libraries were sequenced on a HiSeq2500 (Illumina) to generate 50-base single reads. Mouse testis libraries were sequenced on an Illumina NextSeq 2000 platform using a P2 flow cell with 100 cycles (2 × 50 bp). Mouse heart libraries were sequenced on a NextSeq 2000 platform using a P3 flow cell with 100 cycles (1 × 100 bp). FastQ files were obtained using CASAVA v1.8 software (Illumina).

Quality control of raw sequencing reads was assessed using FastQC v0.11.9 (Andrews, 2010) and aggregated with MultiQC v1.12 (P. Ewels et al., 2016). Adapter trimming and quality filtering were performed using Fastp v0.23.2 (Chen et al., 2018). Reads were aligned to the respective reference genomes using STAR v2.7.10a (Dobin et al., 2013): zebrafish samples were aligned to GRCz11 using Ensembl release 111 (F1 heart) or release 115 (testes), and mouse samples to GRCm39. Gene-level read counts were generated using featureCounts from the Subread package v2.0.3 (Liao et al., 2014). For the mouse F1 heart dataset, transcript-level quantification was additionally performed using Salmon v1.9.0 (Patro et al., 2017) in star_salmon mode. Differential expression analysis was conducted in R v4.4.0 using DESeq2 v1.44.0 (Love et al., 2014) with a design formula of ~condition. Log2 fold change shrinkage was applied using apeglm for mouse datasets and ashr v2.2.63 for zebrafish datasets. Genes with adjusted p-value $< 0.05$ (Benjamini-Hochberg correction) and $|\log2FC| > 1$ (mouse) or $|\log2FC| > 0.58$ (zebrafish) were considered differentially expressed; an additional |log2FC| > 0.58 threshold was applied for volcano plot and pathway analyses in the mouse F1 heart dataset. Batch correction for the zebrafish testes dataset was performed using ComBat-seq from the sva package v3.52.0. For cross-species pathway analysis, zebrafish gene identifiers were converted to mouse (Mus musculus) orthologs using the Alliance of Genome Resources Orthology file (zebrafish testes) or biomaRt v2.60.1 via the Ensembl December 2021 archive (zebrafish F1 heart) (Durinck et al., 2009). Over-representation analysis (ORA) was performed using clusterProfiler v4.12.2 (Wu et al., 2021; Yu et al., 2012) and ReactomePA v1.48.0, with parallel execution via BiocParallel v1.38.0. Gene Set Enrichment Analysis (GSEA) was additionally performed for mouse datasets. Data visualization was performed using ComplexHeatmap v2.20.0, pheatmap v1.0.12 (Kolde, 2019), ggplot2 v3.5.1 (Wickham, 2016), and ggrepel v0.9.5.

**Mass spectrometry data processing and protein identification**

Testes were collected from uninjured males and injured males at 1 and 7 days post-cardiac cryoinjury. Testes from four zebrafish were pooled for each sample, washed in PBS, and snap-frozen. Samples were lysed in 200 µL of 8 M urea, 100 mM Tris/HCl (pH 8.0), and protease inhibitor (Complete, Roche) using 0.4 g Matrix D beads on a FastPrep-24 instrument (animal kidney program; 6.0 m/s, 1 cycle for 40 s). Following centrifugation for 1 min at 16,000 × g at 4°C, the supernatant was transferred to a new vial. The beads were washed once with lysis buffer, and the wash solution was combined with the initial extract.

Proteins were subsequently reduced and alkylated using 10 mM DTT and 50 mM iodoacetate by sequential incubation for 30 min at 37°C. Proteins were precipitated with TCA/acetone (1:8:1 ratio) for 2 h at −20°C. Pellets were washed twice with ice-cold acetone, dried, and stored at −20°C until further processing.

Pellets were reconstituted in 100 µL of 8 M urea and 50 mM Tris/HCl (pH 8.0), and protein concentration was measured using a Bradford assay. An aliquot corresponding to 10 µg of protein was subjected to sequential enzymatic digestion with sequencing-grade LysC for 2 h at 37°C, followed by trypsin digestion overnight at ambient temperature (protease-to-protein ratio, 1:50). Digestion was stopped by addition of TFA to a final concentration of 1%.

For mass spectrometric analysis, 200 ng of peptides were analyzed twice in data-dependent and data-independent modes on a nanoElute 2 timsTOF HT system, as described previously (Ozan et al., 2025). This bottom-up proteomics workflow involved enzymatic digestion of protein extracts into peptides followed by nanoLC-MS/MS analysis.

Mass spectrometry data were searched and quantified using Spectronaut (Biognosys), version 20.2.250922.92449, in directDIA hybrid mode against a concatenated UniProt database and common contaminants. Factory settings were applied with the following adjustments: mass tolerances were set to 20 ppm and 0.05 Th for MS1 and MS2, respectively; precursor q-value, precursor PEP, protein q-value, and protein PEP cutoffs were set to 0.01; and the single-hit protein rule was set to Stratified Single Hit Protein FDR. Single-hit proteins were subsequently excluded from downstream analyses. Search parameters included Acetyl (Protein N-term) and Oxidation (M) as variable modifications, Carbamidomethyl (C) as a fixed modification, and trypsin as the digestion enzyme with a maximum of two missed cleavages. Peptide length was restricted to 7–52 amino acids.

For downstream analysis, intensity-based quantification (IQ) values were calculated using the iq R package (10.32614/CRAN.package.iq) after median normalization and used as the primary abundance measure. Downstream analyses were performed to visualize differential protein abundance using heatmaps and volcano plots.

The mass spectrometry proteomics data have been deposited to the ProteomeXchange Consortium via the PRIDE partner repository with the dataset identifier PXD080667.

**DNA extraction and Whole Genome Bisulfite Sequencing of zebrafish sperm and 1k-cells embryos**

Sperm collected from zebrafish were lysed in lysis buffer (100mM Tris-Cl pH8, 10mM EDTA, 500mM NaCl, 1% SDS, 2% β-Mercaptoethanol) and treated with Proteinase K (20mg/ml) for 2 hours at 55°C.

DNA was then extracted with the QIAamp DNA Mini Kit (Qiagen) following the manufacturer's instructions. 1k-cells embryos were rewarmed gradually on ice and lysed with the AL lysis buffer from the QIAamp DNA Micro kit (Qiagen). Both Proteinase K (Qiagen) and RNase Cocktail (Invitrogen) were used before performing DNA extraction using Phenol:Chloroform:Isoamyl alcohol (25:24:1), followed by ethanol precipitation protocol. Proteinase K-RNAse treatment, as well as the DNA extraction and purification steps, were repeated three times due to the high content of protein and RNA present at this developmental stage.

Purified DNA from both sperm and embryos was quantified by Nanodrop and sent to the Genomics and Epigenetics Division of the Garvan Institute of Medical Research in Sydney, Australia. DNA was further processed for bisulfite conversion. MethylC-seq low-input libraries were prepared as described in (Urich et al., 2015). Briefly, 400ng of genomic DNA was sonicated to an average size of 300 bp using a Covaris sonicator. Sonicated DNA was then purified and end-repaired, followed by the ligation of methylated Illumina TruSeq sequencing adapters. Library amplification (10 PCR cycles) was performed with KAPA HiFi HotStart Uracil+ DNA polymerase (Kapa Biosystems, Woburn, MA). The libraries yield 76-101 million read pairs each. Cytosine non-conversion rate was 0.26-0.3% and CpG sequencing coverage 9.9-13.9x.

Differentially methylated regions (DMRs) between sperm or embryos from uninjured and 7 dpi males were called using the DSS algorithm using the following parameters: minimal length = 100, min number of CpG sites per DMR = 5, p-value threshold = 0.05, delta methylation = 0.1 (10%)). Methylation scores and CpG coverage were visualized on the UCSC genome browser.

**DNA methylation analysis in mouse spermatozoa**

Motile spermatozoa utilized for methylome analysis were cryopreserved. After thawing, sperm were lysed in lysis buffer (1M Tris pH 8, 5M NaCl, 20% SDS, 1M DTT) containing Proteinase K at 55°C for 2 hours and extracted by phenol/chloroform/isoamylalcohol (25:24:1) followed by ethanol precipitation. DNA was resuspended in nuclease-free water to a final concentration of 50ng/µl min. Samples were sent to the Josep Carreras Leukaemia Research Institute Foundation in Barcelona (Spain), where they were bisulfite converted and loaded in the Infinium Mouse Methylation BeadChip (Illumina). Data analysis was performed using the SeSame pipeline in R (Ding et al., 2023; Mani et al., 2022), which sets quality for masked probes, computes p-value using out-of-band probe distribution, and produces Beta methylation scores used for downstream processing (Zhou et al., 2018). The regions detected by Infinium probes were annotated with Sesame using the mouse reference genome (Cavalcante & Sartor, 2017). The differentially methylated genes were subject to GO and pathway enrichment analysis using ClusterProfiler (Yu et al., 2012).

**Sperm nuclei extraction and ATAC-seq**

ATAC-seq of spermatozoa was performed following Ruth Williams' protocol (Williams & Sauka-Spengler, 2021). Shortly, 100’000 spermatozoa were spun down at 500 × g for 5 min at 4°C, washed with 50 µL ice-cold PBS, resuspended in 50 µL lysis buffer (10 mM Tris-HCl, pH 7.4; 10 mM NaCl; 3 mM $MgCl_2$; 0.1% IGEPAL CA-630), centrifuged again, and the supernatant discarded before proceeding to the transposition reaction. The Next Generation Sequencing facility at the University of Bern further performed tagmentation library preparation and sequencing. Tagmentation was performed using the Illumina Tagment DNA Enzyme and Buffer Large Kit (Illumina, 20034198), containing the Tagment DNA Enzyme and TD Buffer. Library amplification was performed using Illumina DNA/RNA UD Indexes Set A, Tagmentation (Illumina, 20091654). Following library preparation, library quality and quantity were assessed using a Qubit 4.0 fluorometer with the Qubit dsDNA HS Assay Kit (Thermo Fisher Scientific, Q32854), an Agilent Fragment Analyzer with the HS NGS Fragment Kit (Agilent, DNF-474), and quantitative PCR using the JetSeq Library Quantification Lo-ROX Kit (Bioline, BIO-68029) according to the manufacturer’s instructions.

Equimolar-pooled libraries were sequenced as 100 bp paired-end reads on an Illumina NovaSeq 6000 instrument using SP and S1 Reagent Kits v1.5 (200 cycles; Illumina, 20040719 and 20028318, respectively). Sequencing run quality was assessed using Illumina Sequencing Analysis Viewer (SAV) version 2.4.7, and base call files were demultiplexed and converted into FASTQ files using Illumina bcl2fastq conversion software version 2.20.

Libraries generated approximately 30-35 million reads per sample. Following quality control and data processing, each sample retained at least 4.6 million and 29 million reads, respectively.

Bioinformatic analysis of ATAC-seq data included trimming of adapters using Trim Galore (Martin, 2011), and alignment of trimmed sequences to the zebrafish reference genome GRCz11 (Ensembl release 115) using Bowtie2 (Langmead & Salzberg, 2012). BAM files were processed and filtered to a high mapping quality threshold of Q30 using SAMtools (Danecek et al., 2021). Duplicates were removed and uniquely mapped reads selected using Picard Tools (Broad Institute, 2019). Tn5 offset correction was applied (+4 bp forward strand, −5 bp reverse strand). Peaks were called on deduplicated BAM files using MACS2 at an FDR threshold of 0.05 (Zhang et al., 2008), yielding a consensus set of 126,797 peaks. Differential accessibility analysis was performed using DiffBind with DESeq2 normalisation (Love et al., 2014; Ross-Innes et al., 2012), comparing ST_ACI 7dpi versus ST_CACI Uninjured samples, with Benjamini–Hochberg correction for multiple testing (padj < 0.05) and log2 fold-change thresholds of 0.5, 1.0, and 2.0. Peak annotation was performed with ChIPseeker (Yu et al., 2015). Pathway enrichment analysis on genes associated with differential peaks was performed using clusterProfiler (Wu et al., 2021; Yu et al., 2012) and ReactomePA (Yu & He, 2016). Motif enrichment analysis was performed on differential peaks using the MEME Suite v5.5.8 (Bailey et al., 2015), specifically XSTREME (Patel & Grant, 2022) against the JASPAR 2022 CORE Vertebrate (Castro-Mondragon & others, 2022), CIS-BP v1.02 (Weirauch & others, 2014) and CIS-BP v2.00 (Lambert & others, 2019) motif databases. BigWig files for visualisation were generated using deepTools (Ramirez et al., 2016), and genomic interval operations were performed with BEDTools (Quinlan & Hall, 2010). The entire preprocessing pipeline was executed via nf-core/atacseq (P. A. Ewels et al., 2020) using Nextflow (Tommaso et al., 2017) within Docker containers.

**Single-nuclei ATAC-seq of adult mouse testes**

Mouse testes were extracted and processed as described in the main Materials and Methods section. Testicular tissue was mechanically minced and enzymatically dissociated using trypsin-EDTA (0.25%), followed by neutralization with DMEM supplemented with FBS (10%). Cell suspensions were passed through a 70 µm cell strainer, subjected to red blood cell lysis (1× RBC lysis buffer), and treated with DNase I (0.1 U/µL final concentration) to remove extracellular DNA and reduce cell clumping. After an additional filtration through a 40 µm cell strainer and cell counting, nuclei were isolated from 1 × 105 - 1 × 106 cells using a detergent-based lysis buffer containing Tween-20 (0.1%), NP-40 (0.1%), digitonin (0.01%), Tris-HCl (10 mM), NaCl (10 mM), $MgCl_2$ (3 mM), and BSA (1%). Nuclei were washed in buffer containing Tris-HCl (10 mM), NaCl (10 mM), $MgCl_2$ (3 mM), Tween-20 (0.1%), and BSA (1%), resuspended in 1× diluted 10x Genomics Nuclei Buffer, and filtered through a 40 µm Flowmi cell strainer. Nuclei were counted and evaluated for structural integrity using a Countess III Cell Counter (Thermo Fisher Scientific). An appropriate volume corresponding to a targeted recovery of 10,000 nuclei was taken for the transposition reaction. Transposed nuclei were subsequently loaded into a Chromium Next GEM Chip H (10x Genomics) and encapsulated into gel beads-in-emulsion (GEMs) using the Chromium Controller (10x Genomics). scATAC-seq libraries were constructed using the Chromium Next GEM Single Cell ATAC Kit v2 (10x Genomics) following the manufacturer's instructions. Library amplification was carried out on a SureCycler 8800 Thermal Cycler (Agilent Technologies). Average fragment size distribution was assessed using a High Sensitivity DNA Kit on a 2100 Bioanalyzer (Agilent Technologies), and library concentrations were quantified using a Qubit Fluorometer (Thermo Fisher Scientific).

Libraries were pooled in equimolar ratios and loaded at a final concentration of 700 pM onto two P3 flow cells (100 cycles) on a NextSeq 2000 System (Illumina) using a paired-end configuration (50 bp Read 1, 8 bp Index 1, 16 bp Index 2, and 50 bp Read 2). Raw sequencing data were demultiplexed, and FASTQ files were generated using the cellranger-atac mkfastq pipeline (10x Genomics).

Raw sequencing reads from mouse testis nuclear samples were processed with Cellranger atac v2.2.0 (10x Genomics, 2019). Each sample was run independently, aligned to the GRCm39/Ensembl 115 reference. Downstream chromatin accessibility analysis was performed in R 4.5.2 using ArchR v1.0.3. Custom genome and gene annotations were derived from the same Ensembl 115 / GRCm39 reference used for alignment. Nuclei were retained if they passed minimum quality thresholds of: TSS enrichment score ≥ 4 and if the Number of unique fragments ≥ 1,000. Doublet scores were computed using ArchR::addDoubletScores() and the predicted doublets were subsequently removed using ArchR::filterDoublets(). Iterative Latent Semantic Indexing (Iterative LSI) was performed on the 500 bp tile matrix (TileMatrix) using ArchR::addIterativeLSI() with two iterations and an initial clustering resolution of 0.2. A UMAP embedding was computed from the LSI dimensions (first 30) using ArchR::addUMAP(). Clusters were identified using ArchR::addClusters() and resolution 1.0, yielding 21 clusters (C1–C21), was selected for all downstream analyses. Gene activity scores were computed from chromatin accessibility within a 2 kb window upstream and 2 kb downstream of each gene's TSS. MAGIC-style imputation weights were added to smooth gene scores across nearest neighbours using ArchR::addImputeWeights(). Marker genes associated with accessibility per broad cell type were identified using ArchR::getMarkerFeatures() on the GeneScoreMatrix.

Cell clustering and annotation were performed to identify the major germ cell and somatic cell populations present in the dataset and to establish the cell type labels used for downstream analyses. Within the somatic compartment, we identified Sertoli cells (*Sox8, Amh, Wt1*), Leydig cells (*Cyp11a1, Cyp17a1, Hsd3b6*), immune cells containing markers for both macrophages (*Apoe, Cd68*) and lymphocytes (*Cd4, Cd8a, Cd3e, Cd79b*), a mixed somatic cell cluster expressing markers associated with epithelial cells (*Krt19*), peritubular myoid cells (*Myh11*), endothelial cells (*Pecam1*), and fibroblasts (*Pdgfra, Col1a2*), and a mixed cell cluster (cluster 2) characterized by markers from multiple testicular cell populations. Among germ cells, we identified spermatogonial stem cells (SSCs; *Cdh1, Sall4, Uchl1*), spermatocytes I (*Piwil2, Sycp3, Meiob, Hormad*), spermatocytes II transitioning into spermatids (with mixed markers), round spermatids (*Piwil1, Acrv1, Spag6*), and elongated spermatids (*Tnp1, Tnp2, Prm1, Prm2*) (Figure 7C). Within spermatocytes I, four distinct clusters were resolved at the main clustering resolution: two preleptotene clusters (*Stra8*), one leptotene–zygotene cluster (*Sycp3*), and one pachytene cluster (*Piwil1*). Based on these annotations, each cluster was assigned both a fine cell type, capturing stage-level resolution (e.g., "Spermatocytes I preleptotene–leptotene"), and a broad cell type, used for all downstream analyses. For the final differential accessibility and motif analyses, we focused on Spermatogonia, Spermatocytes I, Spermatocytes II–differentiating spermatids, Round spermatids, and Elongated spermatids. Group pseudo-bulk coverages were generated per broad cell type using ArchR::addGroupCoverages(). Peaks were called with MACS3 using the ArchR default parameters for single-nucleus ATAC-seq data, and a reproducible consensus peak set was created with ArchR::addReproduciblePeakSet(). A peak-by-cell accessibility matrix was then built using ArchR::addPeakMatrix().

Differentially accessible regions (DARs) between Injured and Uninjured conditions were identified within each broad cell type independently using ArchR::getMarkerFeatures() on the PeakMatrix. The binomial test was applied with bias correction for TSS enrichment and log10(fragment count). Cell types with fewer than 50 nuclei per condition were excluded. Each DAR was annotated with its nearest gene using GenomicRanges::distanceToNearest() against the Ensembl 115 gene annotation. Volcano plots were generated for each cell type using ggplot2. Results were exported as CSV files and BED files for downstream motif analysis. Per-cell-type pseudo-bulk BigWig tracks were exported using ArchR::getGroupBW(). Peaks were visualized for the peaks of interest using Gviz v1.54.0, displaying BigWig data tracks.

Over-Representation Analysis (ORA) and Gene Set Enrichment Analysis (GSEA) were performed in R using clusterProfiler v4.18.4 and ReactomePA. Analysis was run for Gene Ontology categories (Biological Process, Cellular Component, Molecular Function) and Reactome pathways. ORA was applied to DAR nearest genes filtered at FDR $\leq 0.05$ and log2FC $\geq 0.585$. GSEA was applied to the full ranked gene list (ranked by log2FC), using 1,000 permutations. Significant terms were defined by adjusted p-value $< 0.05$ and q-value $< 0.2$. Overlap of DARs across the five spermatogenic cell types (Spermatogonia, Spermatocytes I, Spermatocytes II-differentiating spermatids, Round spermatids, Elongated spermatids) was assessed. GRanges peak coordinates shared within ±100 bp across cell types (GenomicRanges::findOverlaps(maxgap = 100)), visualised as UpSet plots

Per-cell-type DAR BED files (bedtools getfasta) were subjected to motif enrichment analysis using MEME Suite 5.5.8. Random negative control sequences were generated with bedtools random using genome sizes derived from the reference FASTA index. Three motif databases were used CIS-BP 2.00 (Mus musculus), JASPAR 2022 (vertebrates, non-redundant core set), HOCOMOCOv11 (mouse, mono-nucleotide MEME format). XSTREME was run with random genomic background sequences as an explicit negative control (--n), combining *de novo* motif discovery (STREME) with database-matching (SEA). TF overlap across spermatogenic cell types was visualised as Venn diagrams and UpSet plots using ComplexHeatmap (v2.26.1).

For general data wrangling and graph plotting, the following packages were also used: ggplot2 v4.0.2, dplyr v1.2.1, patchwork v1.3.2, rtracklayer v1.70.1, GenomicFeatures v1.62.0, GenomicRanges v1.62.1, BiocParallel v1.44.0, Gviz v1.54.0, ggrepel v0.9.8, and Rsamtools v2.26.0. Analyses were run in a Docker container environment.

#### Movie S1.

Sperm extracted for ATAC-seq. Shown are activated spermatozoa freshly extracted from an adult male zebrafish.
